# Early Radiation Therapy Response Assessment using Multi-scale Photoacoustic Imaging

**DOI:** 10.1101/2025.05.27.656322

**Authors:** Thierry L. Lefebvre, Mariam-Eleni Oraiopoulou, Ellie V. Bunce, Thomas R. Else, Lorna C. Wright, Monika A. Golinska, Lina Hacker, Cara Brodie, Steven Kupczak, Yi Cheng, Lisa Young, Paul W. Sweeney, Sarah E. Bohndiek

## Abstract

There is a critical unmet clinical need to identify biomarkers that predict and detect radiotherapy response in cancer. Using the unique capabilities of multi-scale photoacoustic imaging (PAI) for depicting tumour oxygenation and vasculature *in vivo*, we identified surrogate biomarkers of radiation response in two human breast cancer models (MCF7 and MDA-MB-231), comparing hypofractionated delivery with an ablative single dose scheme. *Ex vivo* immunohistochemistry results underpinned findings from mesoscopic and multispectral tomographic PAI, performed 24h pre-RT, 24h post-RT, and at endpoint. A denser and more mature vasculature of the MCF7 xenografts afforded an improved response to both RT schemes compared to MDA-MB-231, in terms of overall tumour oxygenation, tumour volume and proliferation. Increased intratumoural blood oxygen saturation and oxygen diffusion pre-RT were associated with improved outcomes and decreased proliferation, expected given the oxygen-enhancement effect in RT. *In vivo* PAI revealed the differential effect between ablative courses of RT in both models, with the ablative scheme altering the tumour vasculature as early as 24h post-RT, and pruning the looping vessels and total blood volume at endpoint in the more radiosensitive MCF-7 xenografts. An increase in blood oxygen saturation at endpoint was observed only in the MCF7 xenografts treated with hypofractionated RT, confirming the reduced oxygen consumption of damaged tumour cells, indicative of response. Thus, we showed that PAI could capture early RT response and inform on radioresistance, thus demonstrating promise of PAI as an *in vivo* and future clinical tool to monitor the tumour vascular response to RT.

## Introduction

Hypoxia is a common phenotype in many solid tumours. Uncontrolled cellular proliferation is metabolically demanding, yet the nascent tumour vasculature is often chaotic and poorly supported, leading to inadequate delivery and distribution of oxygen in the growing tumour mass(1). Hypoxia and the associated overexpression of hypoxia-related proteins is frequently associated with tumour aggressiveness, particularly in hormone-dependent cancers such as breast cancer, and poor treatment outcomes, for example, in radiation therapy (RT) (2–4). A particular use-case that intersects these two challenges is the application of neoadjuvant RT in breast cancer treatment, which has shown promise for invasive and locally advanced breast cancer to enable improved resectability and breast-conserving surgery (5–8), and is currently being investigated in clinical trials (ClinicalTrials.gov ID, NCT05479409, NCT05216900, and NCT05412225). In low lineal energy transfer (LET) RT such as X-ray-based therapies, the dose of radiation required to achieve the same biological effect in hypoxic regions is up to 3-fold higher than in normoxic regions (9, 10). Nonetheless, tumour oxygen distribution is typically not assessed or considered clinically during treatment planning even with RT as a first-line treatment in many solid tumours (11), let alone in the neoadjuvant setting.

With the trend towards hypofractionation, *i*.*e*. fewer fractions of higher dose using delivery methods such as stereotactic body/ablative RT (SBRT/SABR), hypoxia becomes a more critical consideration as the tumour microenvironment does not benefit as much from the inter-fraction reoxygenation in conventional RT schemes (12, 13). Real-time visualisation of the spatial distribution of intratumoural hypoxia is necessary to enable dose painting for hypofractionation schemes (14). Furthermore, in ablative courses of RT prescribed with higher dose per fraction (>16 Gy), endothelial cells lining the vasculature will undergo apoptosis and contribute to the tumour response (15–17). Damage to endothelial cells can be mediated through acid sphingomyelinase-dependent and p53-independant membrane alterations, leading to ceramide upregulation and apoptotic signaling, which could acutely exacerbate hypoxia, worsening outcomes specifically through the vascular response to SBRT/SABR (18–20).

The clinical value of imaging in the neoadjuvant RT setting is well established for predicting and assessing treatment response (21, 22); however, it has yet to account for the impact of hypoxia (14, 23, 24). Positron emission tomography (PET) tracers like fluoromisonidazole (^18^FMISO) could be used to map tumour hypoxia (25–27), but adding radiation dose is undesirable for longitudinal imaging. Magnetic resonance imaging (MRI) can determine microcirculatory properties through blood- or tissue-oxygen level dependent signals (28, 29), define sub-volumes for radiation dose escalation based on diffusion (30, 31) and identify perfusion imaging biomarkers (32, 33). Nonetheless, these techniques are constrained by limited spatio-temporal resolution and often require the use of exogenous contrast agents, which comes with costs and potential toxicity concerns (34, 35). Considering the spatio-temporal heterogeneity of hypoxia and angiogenesis in the tumour microenvironment (1, 36, 37), any solution applied in the neoadjuvant setting would need to be low-cost, easy to use at the bedside, and provide high resolution data for immediate interpretation.

In this study, we demonstrate the potential of photoacoustic imaging (PAI) as a solution to these key challenges in radiation oncology (38, 39), demonstrating its value in providing predictive biomarkers, enabling early response monitoring, and visualisation of vascular remodelling. PAI leverages optical contrast for mapping different tissue chromophores at depth by overcoming the optical diffusion limit through ultrasound detection of acoustic relaxation waves. The characteristic high optical absorption contrast of oxy- and deoxy-haemoglobin is ideally suited for visualising perfused vasculature and quantifying tumour blood oxygenation (40, 41). PAI can be applied tomographically at high temporal resolution or at higher spatial resolution with raster-scanning modes (42, 43). Previous studies have shown the ability of PAI to capture RT and chemo-RT response in preclinical models separately using these distinct PAI modes (44–46). However, each examined only part of the roles of the vasculature and tumour oxygenation in the RT response picture. Here, by holistically combining tomographic and mesoscopic PAI, we undertook a longitudinal investigation to establish the potential of quantitative PAI biomarkers to assess vascular morphology and function in two preclinical breast cancer models at 24h before RT treatment, 24h after, and at endpoint (1 week after) with hypofractionated and ablative schemes. Using a thorough immunohistochemistry (IHC) analysis of the models, we demonstrate that PAI captures changes in vascular morphology, including vessel looping structures, alongside intratumoural oxygenation dynamics, which together predict response to treatment, define the early response to treatment and confirming response at endpoint. PAI could therefore play a future role in profiling radiosensitivity, treatment planning and verification, as well as confirming treatment response in the neoadjuvant and adjuvant settings.

## Materials and Methods

A schematic overview of the study design can be found in Figure 1.

**Fig. 1.**
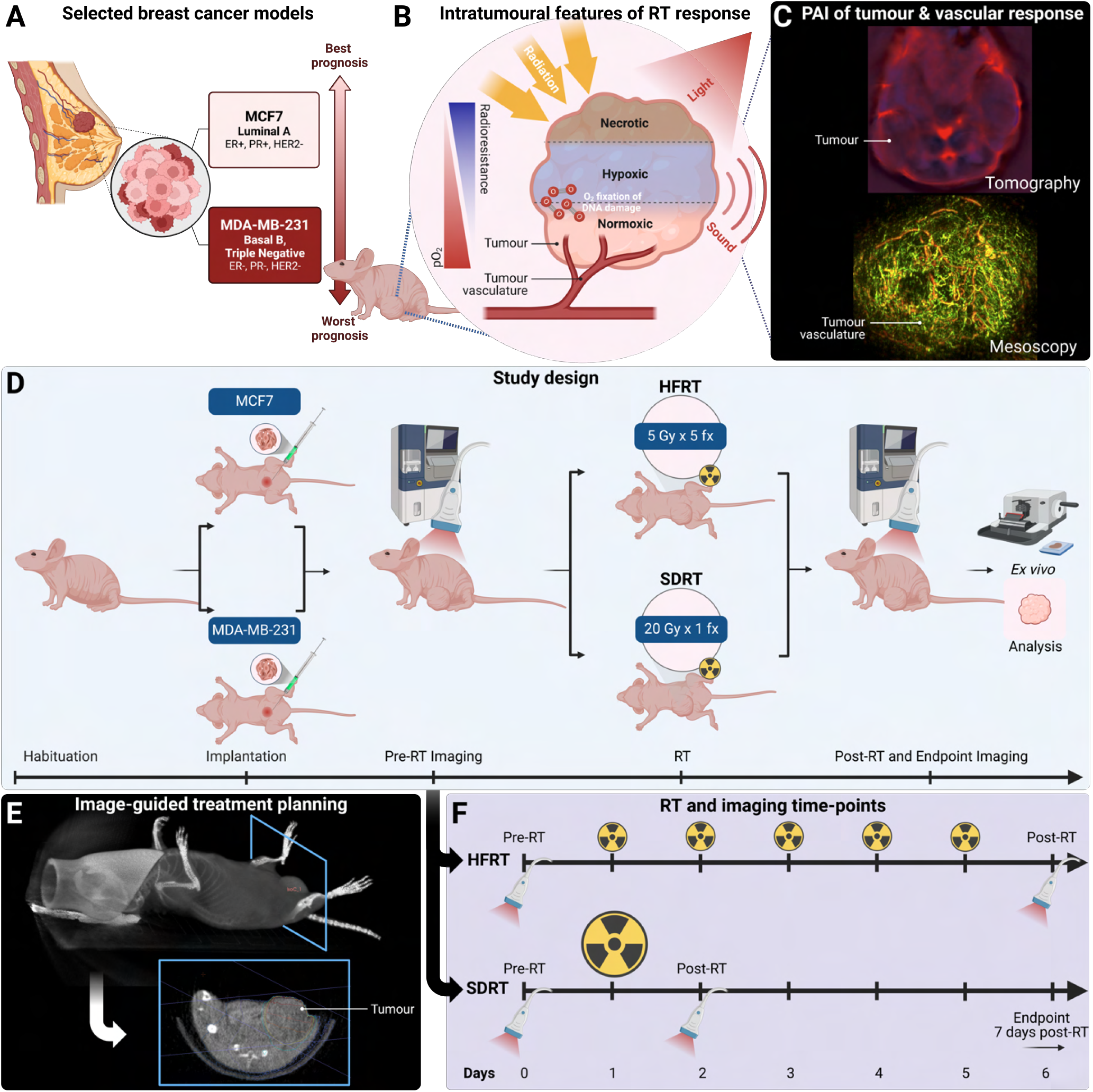
Study Design. A) The human cell lines MCF7, a luminal A subtype ER+ and PR+ breast cancer model, and MDA-MB-231, a triple negative basal B subtype breast cancer model, were selected based on their differences in aggressiveness and radioresistance (47–50). B) Intratumoural features of cancer, such as hypoxia, induce radiation therapy resistance, and the vasculature, being the principal input of molecular oxygen, plays a key role in tumour oxygenation. C) Tumour vasculature can be imaged using photoacoustic imaging, in a tomographic configuration for functional analysis of oxygenation (upper panel), or in a planar raster-scanning configuration at the meso-scale for morphological vascular characterisation (lower panel). D) Study timeline with breast cancer models enrolled into either the control group, or one of the treated groups, *i*.*e*., hypofractionated radiation therapy (HFRT) with 5 Gy per fraction delivered in 5 fraction, or single dose radiation therapy (SDRT) with 20 Gy delivered in a single fraction. E) Treatment simulation on cone-beam computed tomography with a single beam delivered in a moving arc. F) Timeline of imaging and radiation delivery in each treated group, with imaging conducted 24h before, 24h after radiation therapy, and at endpoint (7 days following treatment on average). Created with Biorender.com.

### *In vitro* cell line preparation

Two human breast adenocarcinoma cell lines were selected as part of this study: MCF7, a luminal A subtype with estrogen and progesterone receptors positive, and human epidermal growth factor receptor 2 (HER2) negative (ER+, PR+, HER2-), and MDA-MB-231, a triple negative (ER-, PR-, HER2-) mesenchymal stem-like basal B subtype invasive ductal carinoma (Figure 1A). The models exhibit differential response to RT *in vitro* (survival fraction at 2 Gy of 0.70-0.82 in MDA-MB-231 cell cultures and 0.40-57 in MCF7 cell cultures (47–50)) and when implanted as xenograft models give distinct intratumoural hypoxic and vascular features for *in vivo* PAI (51) (Figure 1B,C). Human cell lines were obtained from the local biorepository of the Cancer Research UK Cambridge Institute at the University of Cambridge, UK. Cells were used only between passages 25-27 (MCF7) or 35-37 (MDA-MB-231). At the start of the study, short tandem repeat (STR) profiling showed 100% match with reference sequences in the two cell lines. Cell lines were passaged and maintained separately in Roswell Park Memorial Institute (RPMI) 1640 Medium (RPMI 1640, Gibco, ThermoFisher Scientific) supplemented with 10% of Foetal Bovine Serum (FBS, Gibco, ThermoFisher Scientific) at 37°C in 5% CO_2_, never exceeding 80% confluence.

### Animal models

Scientific procedures on small animals were performed under the authority of project (PE12C2B96) and personal licenses issued by the Home Office, UK, under the Animals (Scientific Procedures) Act, 1986, and were reviewed for compliance by the local Animal Welfare and Ethical Review Board (compliance form CFSB2226) at the Cancer Research UK Cambridge Institute. All procedures were conducted following the latest guidance on animal welfare (52). Well-being was monitored daily by animal technicians and study conductors. Animals were housed in sterile conditions within hermetic individually ventilated cages with efficiency particulate air filtration, on 12h on/12h off light cycles, and were provided with free access to irradiated sterile food (Mouse Diet 20 Extruded 5R58, PicoLab) and water.

After habituation, subcutaneous cell implantations were performed in the location of the mammary fat pad in eight-week-old immunodeficient female BALB/C nude mice (BALB/c nu/nu, Charles River) (Figure 1D). Cell lines were tested for absence of mycoplasma contamination prior to animal implantation. Inoculations of 1 *×* 10^6^ cells were prepared in a 50 µL solution with a 1:1 ratio of basement membrane extract (Cultrex BME, PathClear, R&D Systems, Bio-Techne) and phosphate-buffered saline (PBS, Gibco, ThermoFisher Scientific), kept on ice. For the ER+ cell line (MCF7) tumour-bearing mice, estrogen pellet implantations (E2-M 17*β*-estradiol 90-day implants, Belma Technologies) were conducted 48h prior to inoculations by surgically implanting a single pellet in the scruff of the neck of animals according to recent protocols (53). When tumours reached a measurable size (> 3 mm in diameter), calliper measurements and animal weighing were performed up to 3 times per week for monitoring tumour growth and animal health. Diameter was measured in the two major orthogonal axes of the growing mass, *a*, the smallest axis, and *b*, the largest axis, and tumour volume, *V*, was calculated as *V* = *ab*^2^*π/*6. Humane endpoints were defined as: i) tumour reaching over 1.5 cm in average diameter; ii) a weight loss surpassing 15% of enrolment weight; iii) acute clinical signs of ill health such as lack of movement, distress, and poor body score; iv) skin conditions such as moist desquamation or bleeding ulceration; and v) in hormone supplemented animals, persistent skin lesions or bladder calcification. During treatment, if signs of weight loss were observed, dietary supplementation in the form of recovery gel (ClearH_2_O DietGel Recovery, Datesand Group) could be provided to animals for their demonstrated value in supporting recovery (54). If hormone supplementation related adverse effects were observed in MCF7-tumour bearing xenografts, animals were closely monitored up to twice a day, applying skin cream (DermaGel) and/or providing bladder massages to facilitate urine passage and limit the potential development of calcification, in consultation with the local named veterinary surgeon.

### Dosimetric quality assurance and treatment delivery optimisation of the preclinical image-guided irradiator

Prior to conducting any *in vivo* experiments, dosimetric quality assurance of the small animal radiation research platform (SARRP, Xstrahl) was performed to ensure that the planned dose corresponded to the actual delivered dose in tissue-equivalent phantoms (solid-water mass density (g/cm^3^), 1.032 *±* 0.005; electron density (*e*^−^/cm^3^), 0.557 *±* 0.001; Sun Nuclear). Dose verification was performed with a National Physical Laboratory-calibrated ionisation chamber (Farmer, PTW Dosimetry) in standard solid-water phantoms in reference conditions (Supplementary Figure 1). The American Association of Physicists in Medicine (AAPM) protocol for 40–300 kV X-ray beam dosimetry was followed (55), using AAPM task group 51 formalism and the measured absolute dose was compared to the reference dose rate of 3.28 Gy/min measured at the initial SARRP commissioning (Supplementary Table 1). It is recommended that differences between prescribed dose and delivered dose be 5% or less (56). Hence, each of the steps including treatment planning should introduce uncertainties under 5% and should add up to a maximum of 5%. In preclinical RT, even though we aim to reduce errors as low as reasonably achievable, higher uncertainties are typically tolerated, with maximal dose output differences of 10% (57). Differences to consider between preclinical and clinical RT are summarised in Supplementary Table 2.

Finally, prior to treatment, fixed beam delivery was compared to moving arc delivery after manually segmenting tumours and surrounding tissue or organs at risk (OAR) as two distinct instances on CBCT images, and resulting dose-volume histograms were assessed (Supplementary Figures 2 and 3). This testing was conducted considering that preclinical irradiators are typically used solely in a top-irradiation fixed set-up. With the improved capabilities of image-guided irradiators, we were provided with the ability to give more conformal treatment and briefly attempted to show the added value of using the SARRP’s abilities through image guidance and moving arc delivery. All dosimetric verification results are provided in the Supplementary Note 1. The implementation of a quality assurance protocol confirmed the suitability of our preclinical RT framework described in the following subsection.

### Radiation therapy planning and delivery

All preclinical image-guided RT experiments were conducted using the small animal radiation research platform (SARRP, Xstrahl) mirroring clinical RT standards (58). RT was delivered under X-ray conebeam computed tomography (CBCT) guidance in the local TG-61-calibrated SARRP with moving arcs optimised to target tumours and spare healthy tissue (55). For imaging guidance, CBCT were reconstructed in commercial treatment planning software (Muriplan, Xstrahl) from acquisitions of 360 projections at one-degree increments in imaging mode (peak energy, 60 kVp; beam filtration, 2.0 mm Al; maximum tube current, 2.8 mA; focal spot size, 0.4 mm; Figure 1E). During treatment planning, a single moving arc spanning 130 degrees was centred on the tumour using CT guidance and the motorised beam collimator size was conformed to the tumour volume (between 7-10mm in both axes). Dose was calculated on the planning software using a superposition–convolution dose engine (59, 60) and delivered on the SARRP using treatment mode at a dose rate of *≈* 2 Gy/min (peak energy, 225 kVp; beam filtration, 0.15 mm Cu; maximum tube current, 13.0 mA; focal spot size, 3.0 mm). Dose distributions were calculated on heterogeneous electron densities obtained from manual intensity thresholding on CBCT, assuming water density for all soft tissues, bone density for bones, and air for regions outside the animal. Source-to-axis distance (centre of the tumour) was fixed and bed positioning and total duration of X-ray delivery were adjusted based on dose calculation from the treatment planning software. Mice were enrolled onto a treatment course of one of the two selected biologically-equivalent RT schemes, when tumours reached a volume of 400 mm^3^: single dose RT (SDRT) of 20 Gy, tested recently in multiple clinical trials in breast cancer patients (61–65), or hypofractionated RT (HFRT) of 25 Gy in 5 fractions of 5 Gy, mirroring the dose fractionation of recent clinical trial in breast cancer patients (66) but in a neoadjuvant context (Figure 1D-F). Tumour-bearing mice from the MCF7 model (*n* = 23) and MDA-MB-231 (*n* = 23) were blindly randomised into control or treated groups (Control, *n* = 13; SDRT, *n* = 18; HFRT, *n* = 15). Enrolled animals treatment and imaging data summaries are provided in Supplementary Table 3.

### *In vivo* photoacoustic imaging

*In vivo* tumour imaging was conducted in anaesthetised mice using mesoscopic (RSOM; Explorer P50, iThera Medical GmbH) and tomographic (MSOT; inVision, iThera Medical GmbH) PAI systems 24h pre-RT, 24h post-RT and at endpoint (mean time after treatment, 7 *±* 3 days) for all mice. Prior to imaging, anaesthesia was induced in animals using 3% isoflurane in a gaseous mix of 50% medical air and 50% pure oxygen. Mice were then transferred to the mesoscopic PAI system on a heated bed kept at 37°C in supine position. Breathing was monitored and maintained between 70-80 breaths per minute, decreasing anaesthetic concentration to 1.5-2% isoflurane after induction, and maintained throughout imaging. The transducer head was manually positioned on the tumour region and coupled to the skin surface using warmed centrifuged ultrasound gel (Aquasonic Clear, Parker Laboratories Inc.). After careful removal of any residual air bubbles, raster-scanning acquisitions were conducted in a 12 × 12 mm^2^ field of view at 20 µm step size, with 3 ns-long laser pulses of 80 mJ in energy at 1 kHz pulse repetition rate, synchronised with ultrasound detection using a single-element transducer with a centre frequency of 50 MHz (*≈* 90% bandwidth). The total scan time for mesoscopic PAI was *≈*7 min.

Anaesthetised animals were then moved to the tomographic PAI system animal holder and a membrane coupled to their skin with warmed ultrasound gel. Prior to acquisition, mice were placed in the tomographic PAI system, then stabilised and acclimatised in the system’s water bath maintained at 36°C for ten minutes. Tomographic acquisitions were performed on the largest tumour cross-section with 6 averages per wavelength (700, 730, 750, 760, 770, 800, 820, 840, 850 and 880 nm). A breathing-gas challenge was performed during tomographic PAI: time-resolved scans were acquired during 5 min on medical air (21% O_2_), after which the breathing gas was switched to pure oxygen (100% O_2_) for another 5 min as described previously (42) (exemplar quantified images in breathing gas challenge in Supplementary Figure 4). The total scan time for tomographic PAI including the gas challenge was *≈*20 min. For pre-RT and post-RT scans, mice were slowly recovered in a heated chamber until active and inquisitive. After the endpoint scan, mice were euthanised with a recognised Schedule 1 method and death was confirmed prior to surgical tumour resection for histopathological processing.

### Photoacoustic image analysis and quantitative imaging biomarkers extraction

All image analyses were conducted in Python 3.9. Motion corrected mesoscopic PAI were reconstructed in 3D using a beam-forming algorithm (67, 68). Images captured the tumour vasculature in a 12 *×* 12 mm^2^ field of view and up to a depth of 4 mm, with in-plan resolution 20 *×* 20 µm_2_ and axial resolution of 4 µm. Mesoscopic photoacoustic images were segmented using the vessel segmentation generative adversarial network (VAN-GAN) (69). Hand-drawn regions of interest (ROI) were selected on *Z*-slices throughout the reconstructed segmented 3D images by an experienced user (LCW), avoiding skin vessels and artefacts, using intensity images for quality assurance verifications. All tumour ROIs were validated by an expert user (TLL). Segmented vascular networks were skeletonised and the total blood volume (BV, µm^3^), the number of perfused vascular connected components (CC, representing the number of sets of vertices connected by path in the vascular skeleton), the vessel segments density (µm^-3^), the average diameters (µm), and the count of looping structures normalised to blood volume (loops, representing a collection of vertices connected by edges in the vascular skeleton forming a path closed on itself, µm^-3^) were extracted as quantitative imaging biomarkers, using previously reported software packages (70, 71) (mesoscopic PAI metrics summary provided in Supplementary Table 4).

Using the Python photoacoustic tomography analysis toolkit (PATATO) (72), averaged PAI tomography images were reconstructed using a back-projection algorithm with in-plane resolution of 75 × 75 µm^2^ at each acquired wavelength. Spectral image series were then downsampled to a resolution of 225 × 225 µm^2^ to increase the signal-to-noise ratio and pixel-wise unmixing was conducted across wavelengths using oxy- and deoxy-haemoglobin absorption spectra. Tumour ROIs were manually selected on the largest tumour cross-section by two experienced users (LCW, TLL). A small ROI around a portion of the large femoral artery was also captured to identify changes in blood oxygenation as a reference. Resulting oxy- and deoxy-haemoglobin parametric maps were quantified in manually delineated tumour and reference ROIs to extract mean total haemoglobin (THb), blood oxygen saturation (sO_2_), and the standard deviation (SD) of sO_2_. We make a note that PAI-derived sO_2_ reported throughout this manuscript is an estimate and not a direct interrogation of true/absolute blood oxygen saturation. Gas challenge tomographic data was analysed using sO_2_ time-series data, identifying the time-point of change to pure oxygen in the reference region selected in the large femoral artery, to quantify and report the change in sO_2_ (ΔsO_2_) under gas challenge in the tumour, and the responding fraction (RF) corresponding to the ratio of voxels with ΔsO_2_ greater than one standard deviation above the baseline (before gas change) mean sO_2_ (tomographic PAI metrics summary provided in Supplementary Table 4).

#### *Ex vivo* histopathological analysis

Excised tumours were processed, and formalin-fixed paraffin-embedded (FFPE) sections were taken for immunohistochemical (IHC) staining and analysis. Briefly, after excision, tumours were fixed in 10% formalin for 24h, and then moved to 70% ethanol for 48h. Resected tumours were then embedded in paraffin, sectioned, and rehydrated. IHC was performed using an automated stainer (BOND, Leica Biosystems) with a bond polymer refine detection kit and 3,3’-diaminobenzadine as a substrate. Tissue sections were stained for endothelial cell marker CD31, smooth muscle cell marker *α*-smooth muscle actin (ASMA), cellular proliferation nuclear marker Ki67, phosphorylated histone nuclear marker of DNA damage *γ*-H2AX, and hypoxia-inducible pro-angiogenic factor marker HIF1-*α* (antibody vendor, dilution and retrieval method provided in Supplementary Table 5). Adjacent 3 µm serial sections were used for CD31 and ASMA staining. Haematoxylin and Eosin (H&E) staining was performed using an automated system (ST5020 Leica, Biosystems). Stained FFPE sections were scanned at 20× magnification using an Aperio AT2 with a resolution of 0.5 × 0.5 µm^2^ (Leica Biosystems).

Analyses were conducted within the whole viable tumour area, excluding skin and necrotic regions, identified on H&E. In HALO (v3.2, Indica Labs), random-forest classifiers were trained by an expert user (C.B.) for distinguishing between the necrotic and viable tissue regions. Classifiers were trained using at least 50 annotations per tissue class per image on a training set of 20-40% of the images and created independently for each stain and then applied across the scanned stained sections. Positive areas of CD31 and ASMA were quantified as a percentage of the total classified tumour area using the area quantification module (v2.4.3). The percentage of positive cells within the classified tumour regions were quantified for Ki67, *γ*-H2AX, and HIF1-*α* with the multiplex IHC module (v3.1.4). To quantify ASMA coverage of CD31-positive tumour vasculature, the CD31 random forest classifier was overlaid onto the ASMA section and its classifier, taking the intersection of the areas as the ASMA vessel coverage, and was reported as a percent of the total viable tumour area (Supplementary Figure 5).

### Statistical analysis

All statistical analyses were conducted using the open-source Pingouin (v0.5.5) and statsmodels (v0.14.3) packages in Python 3.9. Descriptive statistics of the distribution of PAI and IHC biomarkers were computed and visualised with point and bar plots using the seaborn Python package (seaborn, v0.13.2). Unpaired non-parametric two-tailed Mann-Whitney *U* test was used for comparing *in vivo* and *ex vivo* imaging biomarkers between groups and conditions with Holm-Bonferonni multiplicity correction (*m* = 5 comparisons, within tumour model, between control and treated arms, and between controls across models) (73). Bivariate correlations between continuous variables extracted from *in vivo* PAI and *ex vivo* classified IHC regions were assessed with Pearson correlation coefficients, with associated Holm-Bonferonni-adjusted *P*-values. Quantitative imaging biomarkers measured at the different analysed time-points of interest across mice and conditions were analysed as continuous variables with linear mixed effects (LME) models, using as fixed effects the following categorical variables: tumour model (MCF7 or MD-MB-231), treatment scheme (Control, HFRT, or SDRT), and ordinal time-point (pre-RT, post-RT, or endpoint). LME-estimated normalised regression coefficients, standard errors, and Holm-Bonferonni-adjusted *P*-values were reported for each biomarker. Significance level was set at *α* = 0.05 for all adjusted *P*-values.

## Results

### Oestrogen-dependent MCF7 tumours demonstrate greater radiosensivity than triple negative MDA-MB-231 tumours, with improved outcomes using a hypofractionated scheme

As would be expected from the more aggressive triple negative breast cancer model, MDA-MB-231 tumours reached the enrolment volume of 400 mm^3^ faster than MCF-7 (39.3 *±* 6.4 days vs 47.0 *±* 8.2 days, *P*= 0.0058, Figure 2A,B and Supplementary Figure 6) and when left untreated, showed a greater volume increase from enrolment to endpoint compared to MCF7 xenografts (change in volume, 44.01% *±* 13.60% vs. 27.46% *±* 9.29%, *P*= 0.0104; Figure 2C). Following RT, all tumours demonstrated growth inhibition compared to controls across models and RT schemes (*P*<0.001 for all). The largest change in tumour volume between enrolment and endpoint was observed in HFRT-treated MCF7 compared to controls (Figure 2C; −13.65% *±* 13.15% vs. 27.46% *±* 9.29%, *P<* 0.0001), followed by SDRT-treated MCF7 xenografts (3.59% *±* 12.27%, *P* = 0.0006), HFRT-treated MDA-MB-231 compared to untreated controls (12.20% *±* 6.66% vs. 44.01% *±* 13.60%, *P*= 0.00018), and SDRT-treated MDA-MB-231 (9.70% *±* 12.97%, *P* = 0.00036). The most responsive groups also displayed decreased proliferation, as shown by Ki67 on IHC, with significantly lower percentage of Ki67 positive nuclei in HFRT-treated MCF7 xenografts compared to controls (Figure 2D,E; 21.98% *±* 8.37% vs. 50.04% *±* 14.60%, *P*= 0.011), and in the SDRT-treated MCF7 group (23.09% *±* 3.08%, *P* = 0.012). Proliferation in MDA-MB-231 tumours was significantly lower only in the HFRT arm (42.45% *±* 13.32% vs. 23.95% *±* 11.78%, *P*= 0.019). Similarly, DNA damage measured with *γ*-H2AX IHC was significantly increased in all treated groups compared to control mice, across tumour models (Figure 2F,G; *P*<0.01 for all). The measured change in tumour volume was correlated with *ex vivo* IHC markers of response, Ki67 and *γ*-H2AX (*r* = 0.66, *P* = 0.0030; *r* = −0.72, *P* = 0.0008. respectively). Necrotic areas were higher in treated groups than in controls in both models (*P*<0.05, except for SDRT-treated MCF7 vs. MCF7 controls; Supplementary Figure 7). Thus, the oestrogen-dependant MCF7 model was confirmed to be a more radiosensitive breast cancer model than the MDA-MB-231 triple negative breast cancer model. The hypofractionated scheme (HFRT) resulted in improved tumour control in both models, compared to the stereotactic ablative scheme (SDRT), highlighting the benefits of fractionation in RT.

**Fig. 2.**
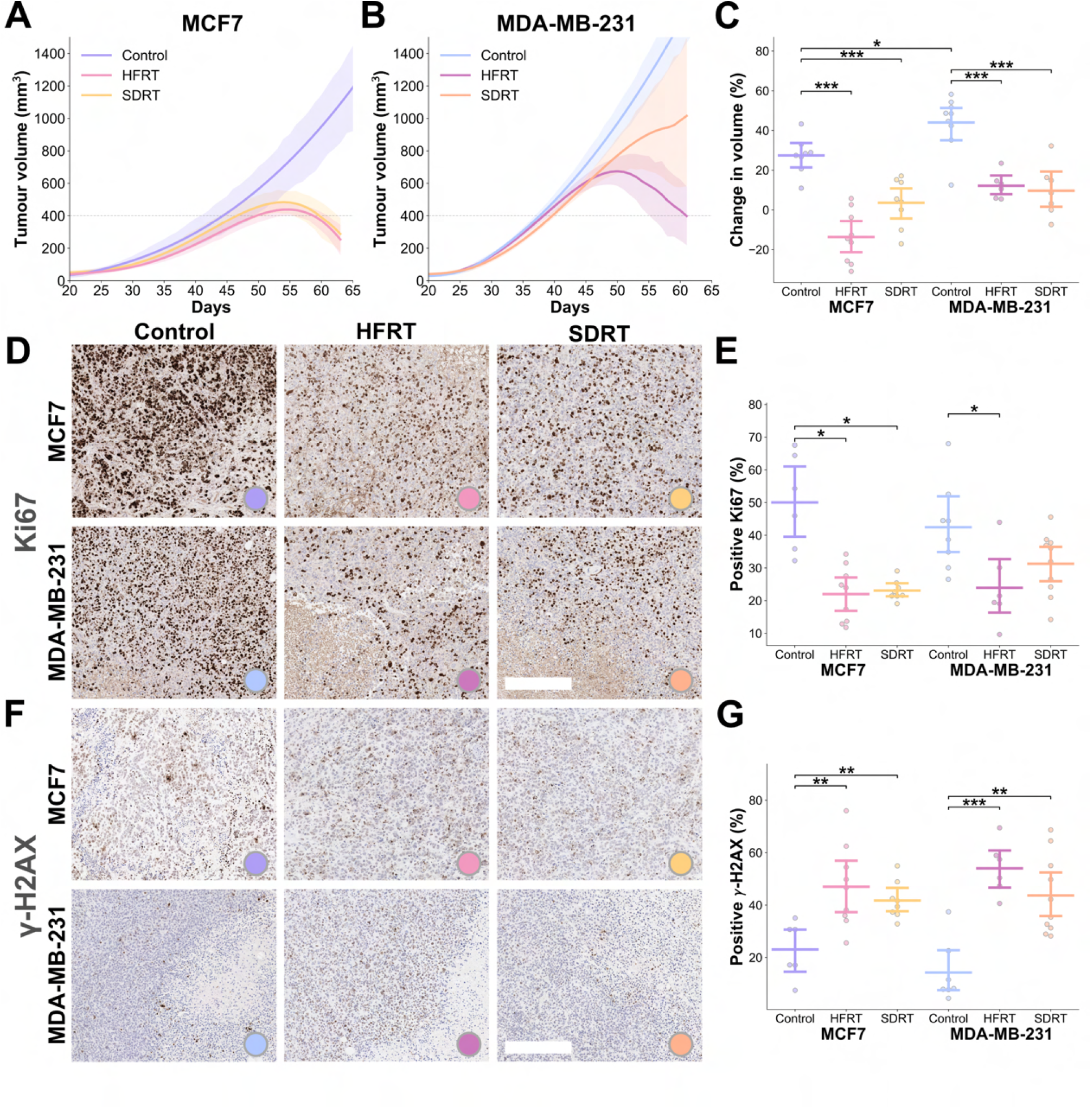
Evaluation of radiotherapy response in the preclinical breast cancer models. Differences in response to selected radiation therapy schemes in the two breast cancer xenografts were assessed with tumour volume and *ex vivo* immunohistochemistry in resected tumours at endpoint. Growth curves assessed by calliper measurements in A) MCF7 and B) MDA-MB-231 models for the untreated (Control), hypofractionated (HFRT)-treated, and single dose (SDRT)-treated groups. Full lines represent average group-wise spline-fitted growth curves, shaded areas are 90% confidence intervals, and dashed lines represent the average enrolment size of 400 mm^3^. C) Percent change in measured tumour volume between enrolment and endpoint (7 days on average post-radiation therapy) demonstrating growth inhibition in all treatment groups. *Ex vivo* immunohistochemistry sections for D) proliferation on Ki67, quantified in E) bar and point plots, and F) DNA damage on *γ*-H2AX, quantified in G) bar and point plots, with bars representing mean and 90% confidence intervals. Scale bar, 200 µm. \**P* < 0.05, \*\**P* < 0.01, \*\*\**P* < 0.001.

To assess the tumour vascular microenvironment in both models, blood vasculature and intratumoural hypoxia were characterised at endpoint on IHC. Double positive CD31-ASMA areas, taken to reflect vascular maturity, were higher in MCF7 than in the MDA-MB-231 tumour sections in the control group (12.39% *±* 1.52% vs. 8.46% *±* 1.83%, *P*= 0.0014; Figure 3A,B). The HIF1-*α* positive area, taken to reflect hypoxia, was also lower in MCF7 xenografts than in MDA-MB-231, but not significantly (13.13% *±* 4.83% vs. 17.28% *±* 3.44%, *P*= 0.11; Figure 3C,D). RT-treated tumours displayed significantly less mature vasculature as seen on CD31-ASMA, compared to controls in both models (*P* < 0.01, except HFRT-treated MDA-MB-321 tumours). HIF1-*α* staining was only significantly decreased in SDRT-treated tumour sections compared to controls in both models (MCF7, *P* = 0.047; and MDA-MB-231, *P* = 0.0045). Hence, MCF7 xenografts were confirmed to possess improved vessel coverage and intratumoural oxygenation in comparison to MDA-MB-231 xenografts, and RT was shown to alter the tumour vascular microenvironment, with greater reduction in vessel maturity and HIF1-*α* staining observed in SDRT-treated tumours.

**Fig. 3.**
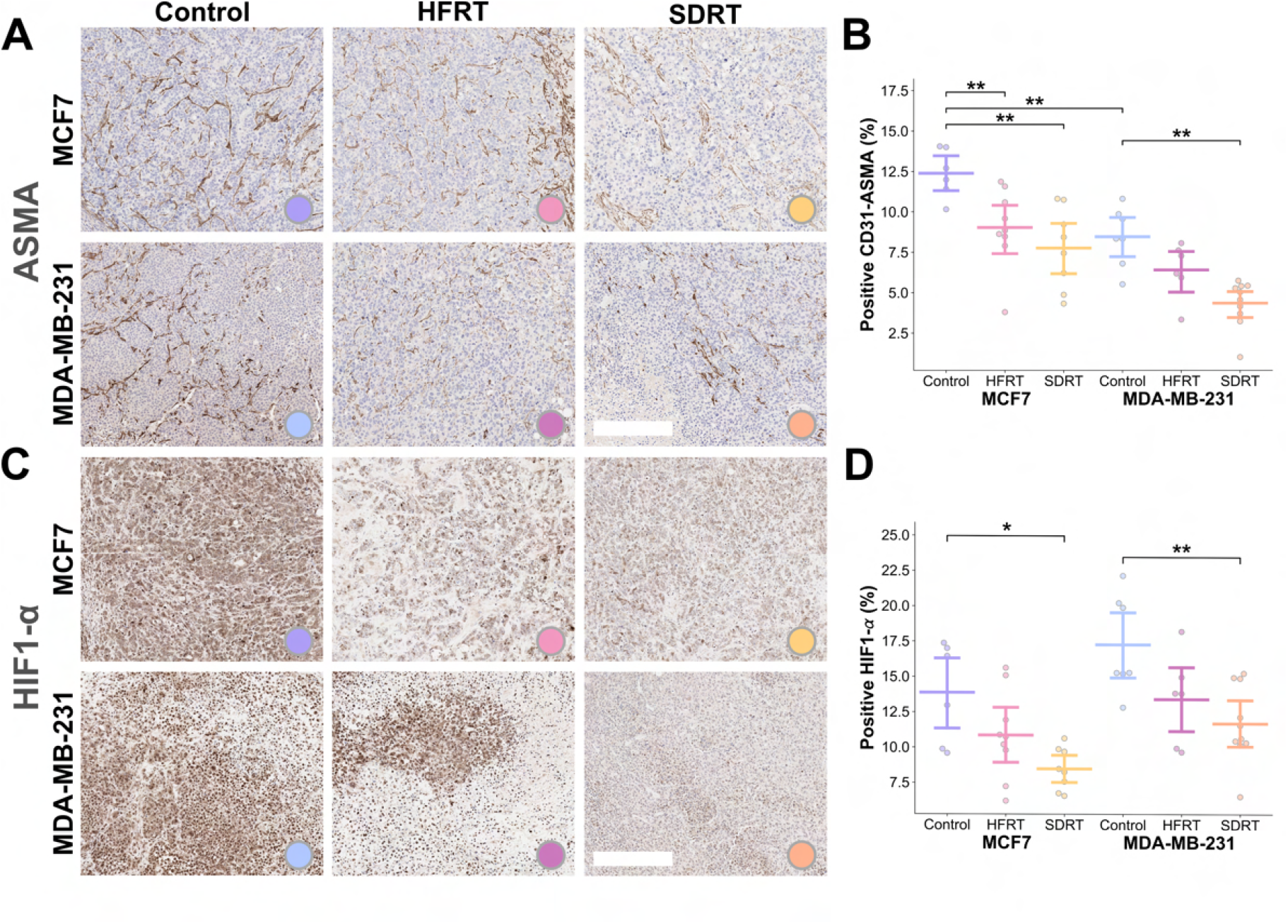
Differences in tumour vasculature and hypoxia after radiation therapy in breast cancer xenografts assessed with *ex vivo* immunohistochemistry. *Ex vivo* immunohistochemistry sections quantified for A) double CD31-ASMA representing mature vasculature coverage (with ASMA sections displayed) in B) bar and point plots, and C) hypoxia on HIF1-*α* in D) bar and point plots, with bars representing mean and 90% confidence intervals. Scale bar, 200 µm. \**P* < 0.05, \*\**P* < 0.01, \*\*\**P* < 0.001.

### Photoacoustic imaging reveals clear differences in vascular microenvironments at baseline in the two models that correlate with radiotherapy outcomes

At enrolment, MCF7 and MDA-MB-231 xenografts were imaged with tomographic PAI 24h pre-RT (Figure 4A). Prior to the administration of any treatment, THb was higher in MCF7 tumours than MDA-MB-231 tumours (0.126 *±* 0.026 vs. 0.102 *±* 0.020, *P*= 0.0032; Figure 4B) as was sO_2_, although non-significantly (0.408 *±* 0.096 vs. 0.362 *±* 0.055, *P*= 0.050; Figure 4C), in line with previous findings (51). Intratumoural distribution of sO_2_ displayed less heterogeneity in the MCF7 compared to the MDA-MB-231 model (SD of sO_2_ pre-RT, 0.091 *±* 0.035 vs. 0.137 *±* 0.050, *P*= 0.047; Figure 4D). Upon administration of a change in breathing gas to 100% oxygen, the change in sO_2_ and the fraction of pixels responding to gas challenge were both higher in the MCF7 tumours compared to the MDA-MB-231 (ΔsO_2_ pre-RT, 0.043 *±* 0.014 vs. 0.034 *±* 0.013, *P*= 0.039; RF pre-RT, 0.588 *±* 0.188 vs. 0.469 *±* 0.097, *P*= 0.018 Figure 4E,F) indicating greater vascular maturity.

**Fig. 4.**
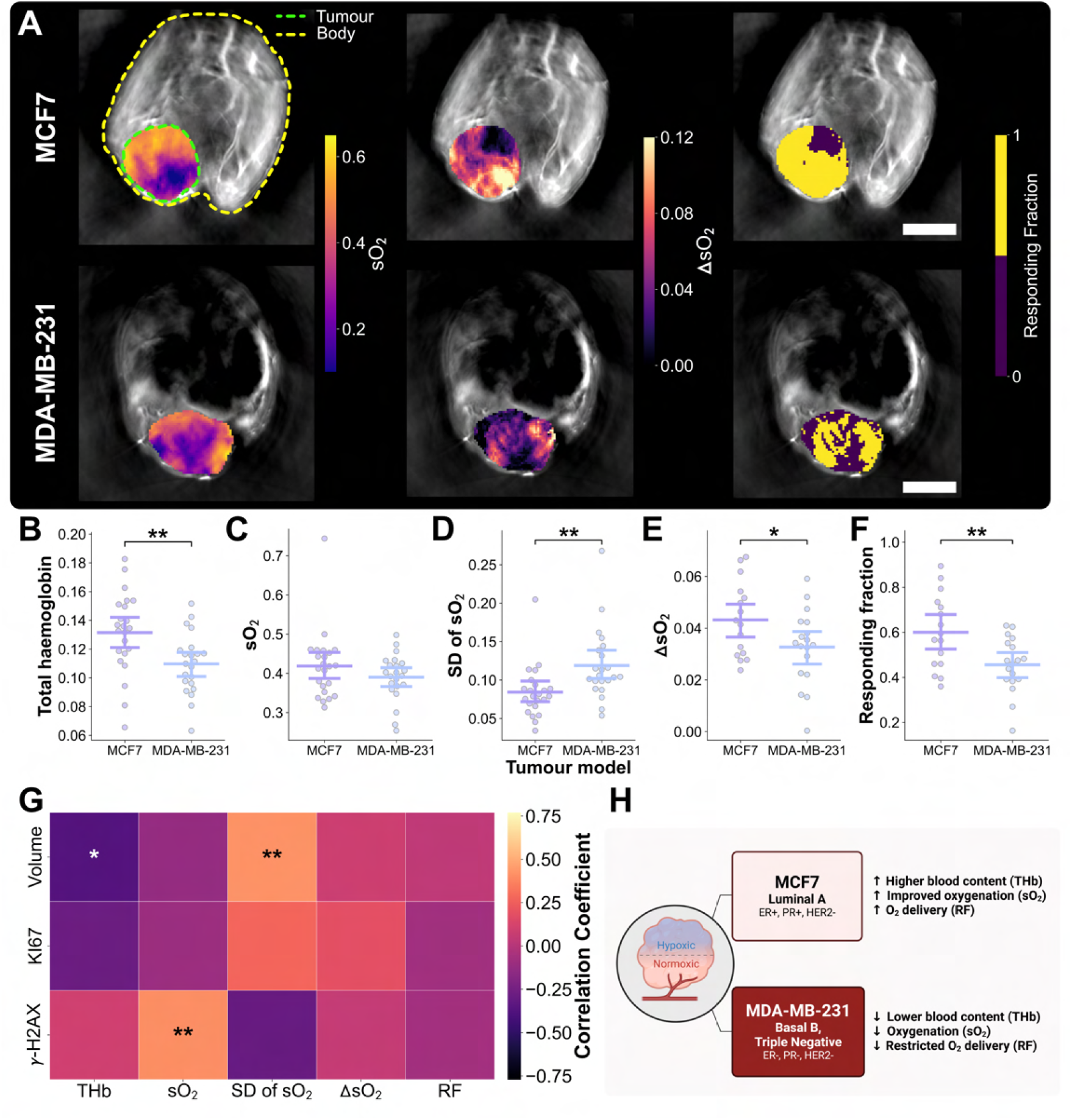
Tomographic PAI delineates models based on intratumoural blood content and oxygenation at baseline. A) Exemplar MCF7 and MDA-MB-231 tumour-bearing mice imaged at baseline (enrolment) with multispectral tomographic PAI, overlaid with quantitative parametric maps (from left to right) of sO_2_, and in response to a breathing gas challenge, ΔsO_2_ and responding fraction. Scale bars, 2 mm. Bar plots of B) total haemoglobin, C) blood oxygen saturation (sO_2_) D) spatial standard deviation (SD) of sO_2_, E) ΔsO_2_ under gas challenge, and F) responding fraction where each point represents an independent tumour. G) Correlation heatmap of *in vivo* tomographic PAI quantitative biomarkers and *ex vivo* immunohistochemistry parameters. H) Summary schematic of vascular microenvironment assessment using baseline tomographic PAI biomarkers in both models. \**P* < 0.05, \*\**P* < 0.01, \*\*\**P* < 0.001. Created with BioRender.com.

We evaluated correlations across both models between pre-RT PAI biomarkers and endpoint histopathological IHC markers to assess the potential of tomographic PAI to reflect the underlying molecular changes that could predict future RT response. THb and SD of sO_2_ at baseline were correlated with the change in tumour volume (*r* = −0.38, *P* = 0.016, and *r* = 0.42, *P* = 0.0041, respectively; Figure 4G). Baseline sO_2_ also showed a significant correlation with *γ*-H2AX (*r* = 0.41, *P* = 0.0059; Figure 4G). These findings indicate that increased blood content pre-RT and lower heterogeneity in oxygenation were associated with improved RT efficacy. Higher tissue oxygenation was related to a higher extent of DNA damage, which would be expected given the well-known oxygen enhancement effect. Overall, using tomographic PAI, MDA-MB-231 xenograft tumours displayed higher tissue oxygenation heterogeneity, lower tumour blood content, and weaker intratumoural oxygen diffusion ability than the MCF7 model, which correlated with poorer treatment response at endpoint (Figure 4H).

Considering the vascular architecturein more detail, mesoscopic PAI indicated a denser peripheral perfused vasculature in MCF7 xenografts compared to MDA-MB-231 xenografts (vascular density, 236.4 *±* 84.5 µm^*−*3^ vs. 153.9 *±* 41.2 µm^*−*3^, *P*= 0.00006; Figure 5A,B). A higher number of vascular connected components were also observed in MCF7 tumours, corresponding to the number of perfused vessel segments connected in the captured vascular skeletons (103.13 *±* 57.48 vs. 64.09 *±* 20.61, *P* = 0.00185; Figure 5C). Although the density of vascular networks between models was different, the overall vascular morphology (BV, diameters and looping structures, Supplementary Figure 8) captured close to the tumour surface was not significantly different. The measured pre-RT blood volume (BV) was significantly correlated with biomarkers of RT response at endpoint (change in tumour volume *r* = −0.34, *P* = 0.022, Ki67 *r* = −0.48, *P* = 0.0016, *γ*-H2AX *r* = 0.50, *P* = 0.0017; Figure 5D). Similarly, increased loops normalised to BV was associated with decreased proliferation (Ki67, *r* = −0.49, *P* = 0.0018) and increased DNA damage (*r* = 0.36, *P* = 0.034). Overall, MCF7 xenografts displayed a denser perfused vascular network at tumour periphery compared to that of the MDA-MB-231, and measured BV and looping structures were associated with improved tumour response to RT (Figure 5E).

**Fig. 5.**
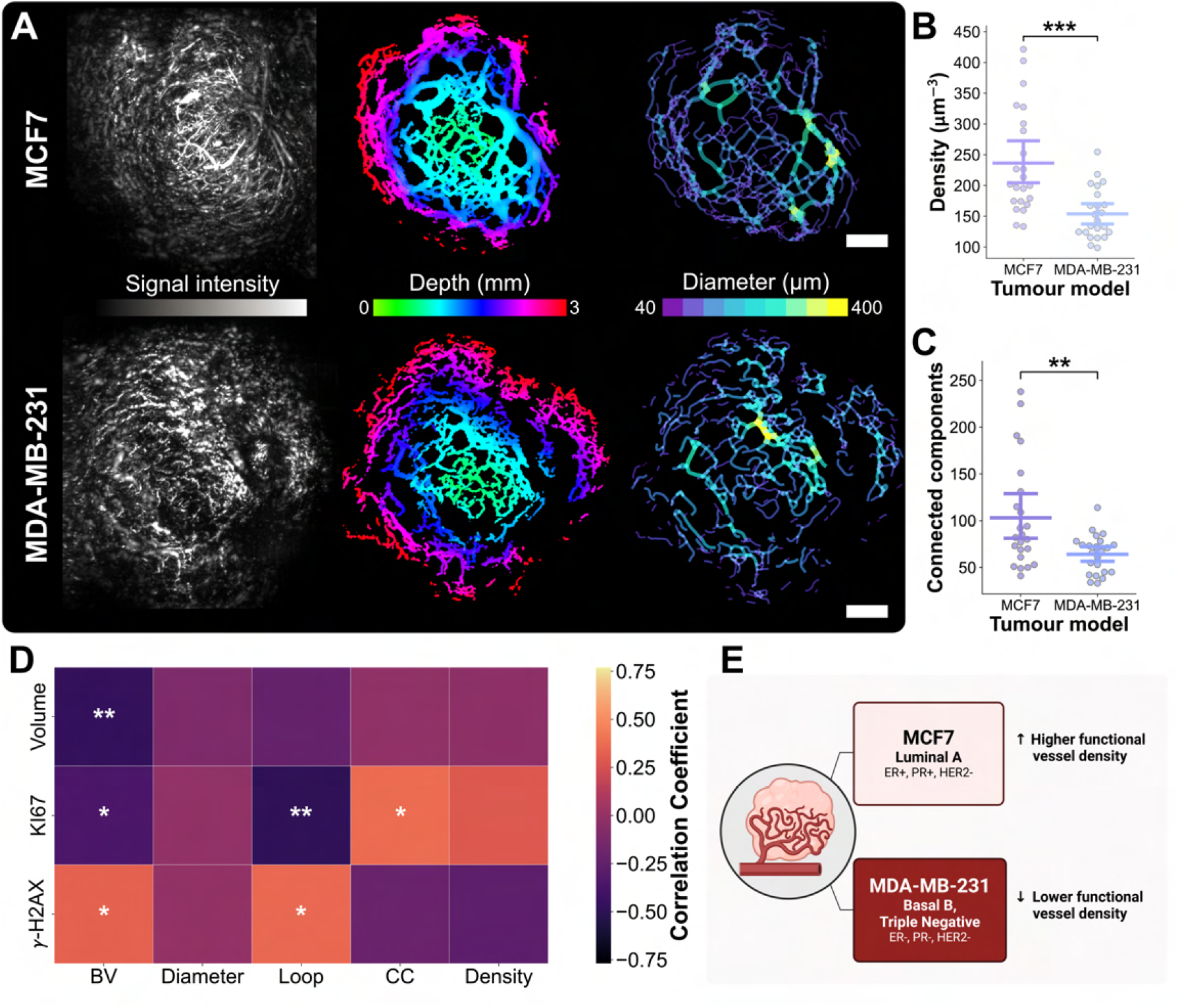
Mesoscopic PAI detects underpinning differences in vascular density and architecture. A) MCF7 and MDA-MB-231 tumour-bearing mice imaged at baseline (enrolment) showing (from left to right) mesoscopic PAI signal intensity, depth-resolved segmented vasculature and a skeletonised diameter map. Scale bars, 2 mm. Bar plots of B) mesoscopic PAI vascular density and C) number of connected components across the vascular networks. D) Correlation heatmap of *in vivo* mesoscopic PAI quantitative biomarkers and *ex vivo* immunohistochemistry parameters. E) Summary schematic of vascular microenvironment assessment using baseline mesoscopic PAI biomarkers in both models. \**P* < 0.05, \*\**P* < 0.01, \*\*\**P* < 0.001. Created with BioRender.com.

### Longitudinal photoacoustic imaging can predict and detect early tumour response to radiotherapy

PAI tomography scans were acquired longitudinally including at 24h post-RT in both treatment arms (Figure 6A,B). On tomographic PAI, the change in THb from baseline across time-points, tumour models and treatment arms was explained with LME modelling. On average, THb increased over time-points, in groups treated with SDRT, and differed between models (time-point normalised regression coefficient [NRC], 0.069 *±* 0.026, *P* = 0.009; SDRT arm NRC, 0.159 *±* 0.065, *P* = 0.0014; and tumour model NRC, 0.119 *±* 0.053, *P* = 0.025, respectively; Figure 6C,E). In fact, when comparing between treatment arms in both tumour models, 24h post-RT THb values normalised to pre-RT scan were greater in SDRT-treated groups compared to controls, although not significantly.

**Fig. 6.**
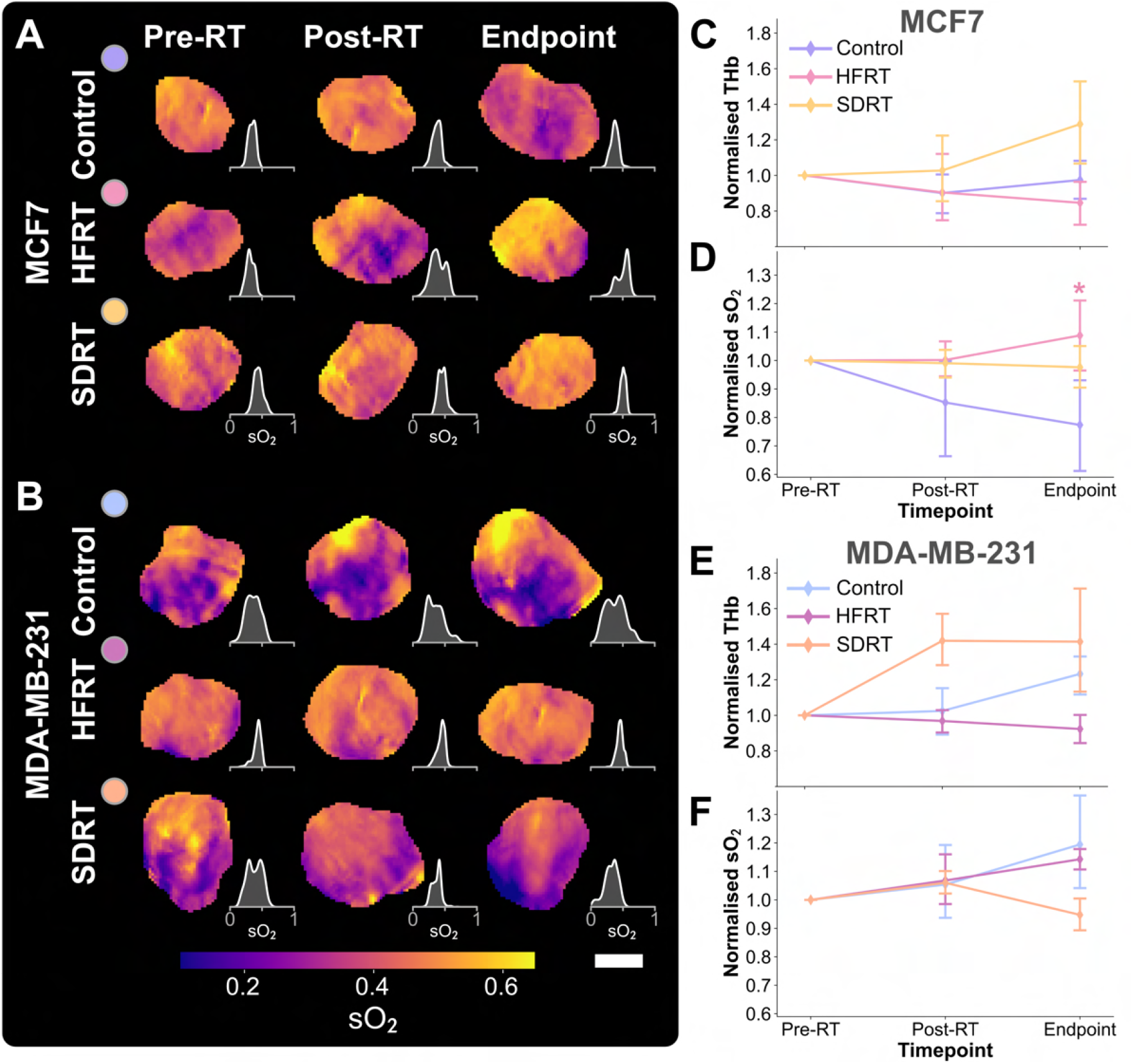
Longitudinal *in vivo* tomographic photoacoustic imaging detects radiotherapy response in breast cancer models. Exemplar blood oxygen saturation tumour maps with paired density plots in A) MCF7 and B) MDA-MB-231 xenografts, across time-points and treatment groups. Scale bar, 2 mm. Tomographic photoacoustic imaging biomarkers normalised to pre-treatment scan in MCF7 and MDA-MB-231 tumour-bearing mice, respectively, C,D) total haemoglobin (THb), and E,F) blood oxygen saturation (sO_2_). \**P* < 0.05, \*\**P* < 0.01, \*\*\**P* < 0.001.

LME modelling revealed that sO_2_ changed modestly in our dataset, and only significantly from baseline between the different tumour models (tumour model NRC, 0.071 *±* 0.036, *P* = 0.046; Figure 6D,F). No significant sO_2_ changes were measured 24h post-RT. Nonetheless, sO_2_ quantified on the 24h post-RT scans was indicative of treatment response across groups when conducting bivariate correlation analysis with endpoint IHC markers (Supplementary Figure 9). sO_2_ at 24h post-RT also showed strong correlation with change in tumour volume, proliferation, and DNA damage (change in tumour volume, *r* = −0.47, *P* = 0.014; Ki67, *r* = −0.55, *P* = 0.0053; and *γ*-H2AX, *r* = 0.54, *P* = 0.0051). Moreover, mean sO_2_ negatively correlated with endpoint hypoxia (HIF1-*α, r* = −0.42, *P* = 0.0188), while its SD was also predictive of endpoint decrease in tumour size (*r* = 0.46, *P* = 0.034). SD of sO_2_ can be appreciated in the sO_2_ density distribution subplots of Figure 6A,B. Thus, PAI tomography could capture early increases in intratumoural blood content, which are likely related to radiation-induced transient inflammatory activity following high doses of radiation, and longitudinal oxygenation mapping was found to be indicative of endpoint tumour control.

On mesoscopic PAI, blood volume decreased most in SDRT-treated mice, then in HFRT-treated groups, and was altered with increasing time-points according to LME analysis (SDRT arm NRC, −0.500 *±* 0.144, *P* = 0.001; HFRT arm NRC, −0.405 *±* 0.150, *P* = 0.007; and time-points NRC, 0.111 *±* 0.055, *P* = 0.045; Figure 7A,B). Similarly, vascular density and connected components changed most over time, as shown by LME modelling (time-points NRC, 0.082 *±* 0.034, *P* = 0.015; 0.188 *±* 0.050, *P* = 0.0002; SDRT arm NRC, −0.373 *±* 0.143, *P* = 0.009; and HFRT arm NRC, −0.326 *±* 0.149, *P* = 0.029). Normalised looping structures were decreased in both treatment groups and changed with time-points (HFRT arm NRC, −0.384 *±* 0.152, *P* = 0.012; SDRT arm NRC, −0.373 *±* 0.146, *P* = 0.011; and time-point NRC, 0.134 *±* 0.054, *P* = 0.014). Average diameters extracted from longitudinal mesoscopic PAI did not reveal any difference between models, treatments or time-points.

**Fig. 7.**
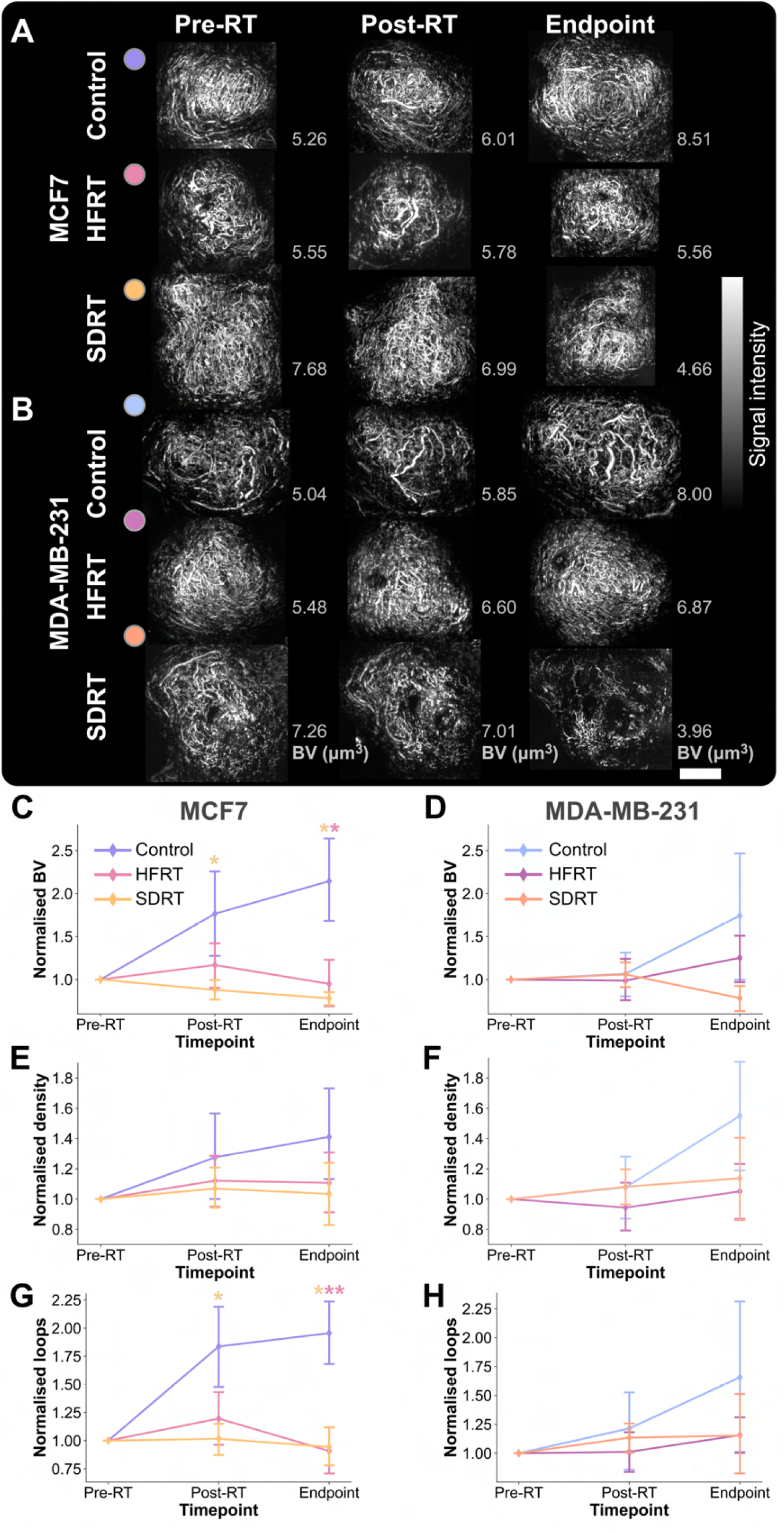
Longitudinal *in vivo* mesoscopic photoacoustic imaging of treatment response in breast cancer models. Exemplar maximal intensity projections of vascular networks on photoacoustic mesoscopy in A) MCF7 and B) MDA-MB-231 xenografts, across time-points and treatment groups. Scale bar, 4 mm. Mesoscopic photoacoustic imaging biomarkers normalised to pre-treatment scan in MCF7 and MDA-MB-231 tumour-bearing mice, respectively, C,D) blood volume (BV), E,F) vascular density, and G,H) looping structures. \**P* < 0.05, \*\**P* < 0.01, \*\*\**P* < 0.001.

Blood volume was decreased in the MCF7 xenograft model within 24h post-RT in SDRT-treated mice, before any change in tumour size was observed (normalised BV, *P* = 0.020; Figure 7A), but not in MDA-MB-231 (Figure 7B). At this early time-point, the vascular density was also decreased in both models in treated groups, but not significantly (Figure 7C,D). In the more radiosensitive MCF7 model, the number of looping structures was significantly decreased compared to control in SDRT-treated mice (*P* = 0.029; Figure 7E), thus decreasing the overall tortuosity of the vascular network, which was not observed in the more radioresistant MDA-MB-231 xenografts (Figure 7F). The observation of significant BV differences only in MCF7 tumours treated with ablative doses of radiation likely indicates endothelial cell damage at high doses. Finally, the measured BV 24h post-RT was predictive of Ki67 positive areas at endpoint (*r* = 0.32, *P* = 0.0384; Supplementary Figure 9), and BV and loops (normalised to BV) were both correlated with hypoxia (HIF1-*α, r* = −0.34, *P* = 0.0487; and *r* = −0.58, *P* = 0.00013; respectively). Thus, PAI mesoscopy provided early detection of response in the tumour vasculature, prior to any change in tumour size, indicating its sensitivity to radiation-induced vessel pruning in the radiosensitive model.

### Differential vascular response to radiation therapy schemes revealed by photoacoustic imaging at endpoint

At endpoint, seven days on average following the last day of treatment, growth was significantly delayed in treated groups of both investigated breast cancer models (Figure 2). A final imaging session was conducted prior to tumour excision for *ex vivo* analysis. THb increased between pre-RT and endpoint in SDRT-treated MCF7 and MDA-MB-231 xenografts compared to their respective controls, and while the opposite trend was observed with HFRT-treated groups (Figure 6C,D), neither reached statistical significance. sO_2_ was higher in treated MCF7 tumours compared to controls, but only significantly in HFRT-treated mice (*P*= 0.017), while no significant difference was observed in the more radioresistant MDA-MB-231 model. In terms of imaging-IHC correlations, THb was negatively correlated with hypoxia (HIF1-*α, r* = −0.45, *P* = 0.038).

On mesoscopic PAI, the change in BV from baseline was significantly lower in MCF7 tumour-bearing treated groups compared to controls (HFRT, *P* = 0.028; and SDRT, *P* = 0.017; Figure 7A), and was also lower in treated MDA-MB-231 xenografts but not significantly (HFRT, *P* = 0.35; and SDRT, *P* = 0.082; Figure 7B). The same trend was observed in measured blood vessel density in both MCF7 and MDA-MB-231 xenografts, but was not significant. The normalised count of looping structures was significantly reduced at endpoint in treated MCF7 tumours only (HFRT, *P* = 0.0079; SDRT, *P* = 0.014). Interestingly, the group averages of total BV change from baseline decreased in the same fashion as the CD31-ASMA percent area in both models, with higher quantified values in control, HFRT and then SDRT (Figure 3A,B). BV was also negatively correlated with the change in tumour size (*r* = 0.53, *P* = 0.0042; Supplementary Figure 9). CC and density at endpoint were both positively correlated with proliferation assessed on Ki67 IHC (*r* = 0.48, *P* = 0.0052; and *r* = 0.42, *P* = 0.023; respectively) and CC were also negatively correlated with the extent of DNA damage (*r* = −0.42, *P* = 0.035). Taken together, our findings indicate a higher volume of denser functional blood vessels at endpoint is correlated with poorer overall tumour response, or radioresistance, in the investigated breast cancer models.

## Discussion

The challenge of treating hypoxic solid tumours with radiotherapy has long been recognised, yet is rarely acted upon in treatment practice. With the increasing use of hypofractionation, hypoxia becomes a more critical consideration and there is a clear need to develop suitable imaging biomarkers to detect and monitor changes in tumour vascular architecture and function in response to RT treatment. In this study, following implementation of dosimetric quality assurance protocols (74), we sought to demonstrate the potential of PAI in this context, comparing hypofractionated and ablative single dose RT schemes in breast cancer xenografts exhibiting differential radiation responses showing wide-ranging potential for PAI guidance of RT (Figure 8).

**Fig. 8.**
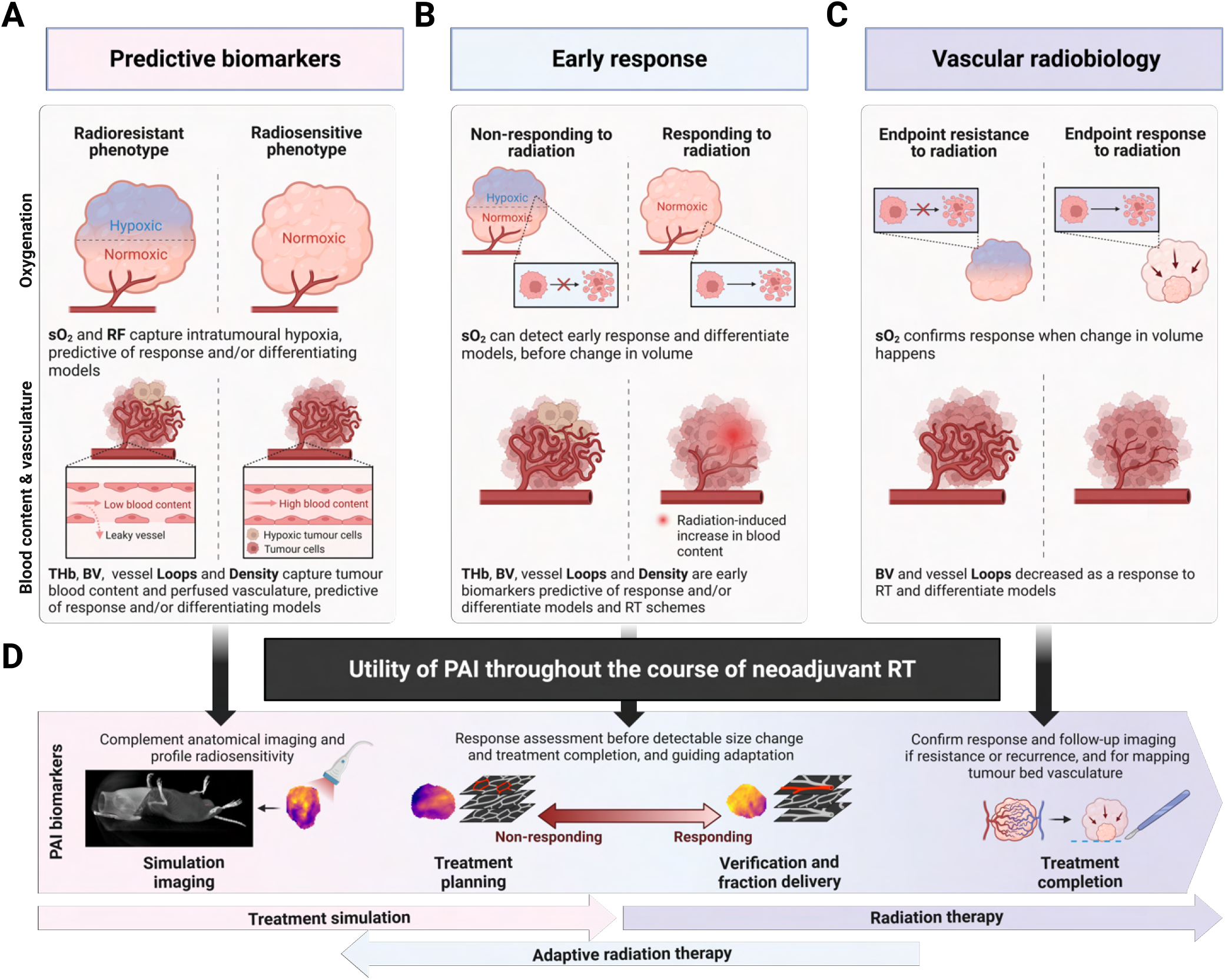
Conceptual schematic of reported PAI biomarkers for radiation response assessment in the tumour vascular microenvironment of breast cancer models. A) PAI biomarkers 24h pre-RT are predictive of response in terms of endpoint change in volume, DNA damage, and proliferation; B) and remain predictive and identify early response 24h post-RT. C) Endpoint imaging reveals the vascular response to RT with endothelial cell apoptosis leading to microvascular disruption, identified as a decreased loop count and overall blood volume. D) Potential use of PAI within the neoadjuvant RT framework, at baseline for profiling tumour radiosensitivity, for mapping intratumoural oxygenation to enable dose painting and further adaptation over the course of treatment, the ability to detect and predict response, and value in confirming response following treatment completion for improved tumour resectability guidance. Note: sO2, blood oxygen saturation; RF, responding fraction; THb, total haemoglobin; BV, blood volume; PAI, photoacoustic imaging; RT, radiation therapy. Created with BioRender.com.

*Ex vivo* analysis of the two models aligned with the literature, showing MCF-7 tumours to have a more developed and mature vasculature, associated with lower hypoxia (51, 75) and radiosensitivity (47–50, 76). Both models showed gradual reductions in vessel maturity from control, to HFRT and then SDRT arms, alongside a characteristic decrease in hypoxia through HIF1-*α* IHC in SDRT-treated groups.

*In vivo* imaging biomarkers evaluated in this study revealed pre-RT differences between the MCF7 and MDA-MB-231 models in terms of tumour blood content, oxygenation, tissue oxygen diffusion, and vascular morphology, aligned with prior work (51). We found that increased THb and decreased heterogeneity of oxygen saturation (SD of sO_2_) prior to treatment were associated with tumour control at endpoint across both models. It is worth noting that although MDA-MB-231 have been reported to have a higher microvascular density than MCF7 on IHC (51, 75), the mesoscopic PAI measurements demonstrated a significantly denser network in the MCF7 model, even though the total blood volume did not differ. This discrepancy could be explained by the fact that mesoscopic PAI depicts only the blood-filled vessels larger than its in-plane resolution (*≈* 20 µm) at the peripheral rim of the tumour, whereas IHC analyses the core and samples all vessels expressing endothelial cell proteins. Of note, a previous study has shown that orthotopic breast cancer tumours used in our study produce more vascularised tumours compared to subcutaneous tumours, used in prior studies (77). Encouragingly though, increased blood volume pre-RT in mesoscopic PAI correlated with endpoint response, similar to whole tumour-evaluated THb on tomographic PAI.

Others have reported tumour blood oxygen saturation (sO_2_) pre-treatment as a predictive biomarker for RT response in patient-derived xenografts of head & neck cancer or pancreatic cancer models (44, 78, 79). However, even though sO_2_ was higher on average in the more radiosensitive MCF7 at baseline, we did not observe a significant difference in our study; sO_2_ was only significantly different at endpoint. Most interestingly, we were able to assess *in vivo* the well-known oxygen-enhancement effect in RT (10), by finding pre-RT a correlation between sO_2_ as measured with PAI tomography and the extent of DNA damage at endpoint, which was consistent across models and treatment groups at the individual level. Our results confirm those of a previous study using nanosonophores as a PAI contrast agent for oxygen sensing, which also showed that pre-RT oxygen distribution was associated with post-RT DNA damage *ex vivo* on *γ*-H2AX in breast cancer patient-derived xenografts (80).

Change in blood oxygen saturation and the responding fraction under gas challenge pre-RT were were both significantly higher at baseline in the MCF7 groups than in the MDA-MB-231, indicating better intratumoural oxygen diffusion, linked to higher radiosensitivity in MCF7. A prior study investigating the same models revealed that blood oxygen level dependent-MRI ΔR2*, a surrogate marker for oxygen delivery under gas challenge, showed improved blood oxygen diffusion in MCF7 compared to MDA-MB-231 xenografts, in-line with our findings (75). Increased tumour ΔsO_2_ pre-treatment was also demonstrated to be a predictive biomarker in pancreatic cancer models (79, 81), highlighting the potential of using a breathing gas challenge to extract predictive PAI biomarkers.

Importantly, PAI revealed the differential effect of ablative (SDRT) and hypofractionated (HFRT) courses of RT in the two models. Since RT has the potential to stimulate tumoural neovascularisation and perfusion through an acute inflammatory response (82), and RT can also prune hypoperfused vessels (71, 83–85), blood flow to the tumour may be transiently increased following a dose of RT, thereby increasing THb. Just 24h after radiation delivery, an increase in tumour blood content was observed in SDRT-treated mice, indicating a direct vascular contribution to radiation response. We also confirmed that vessel loops decrease significantly at both 24h after treatment and at endpoint, demonstrating vessel pruning. Considering oxygenation response, we that higher sO_2_ 24h post-RT was highly correlated with endpoint tumour control across groups and sO_2_ was significantly increased at endpoint in the more responsive MCF7 tumours. Increased sO_2_ post-SDRT has also been observed by others (46, 86, 87). Overall, improvement in oxygenation can likely be attributed to the reduced oxygen consumption of damaged tumour cells, indicative of response, rather than to an increase in oxygen supply or oxygen diffusion.

Despite these promising findings, there are limitations to our study. A first key limitation is that the imaging modalities investigated here only provide surrogate biomarkers of complex biological processes focusing on tissue vascularisation, blood content and oxygenation – and not cell death – and must therefore be carefully interpreted. For instance, tomographic PAI-derived blood oxygen saturation is not an absolute oxygenation measurement, but provides insightful relative sampling of the intratumoural oxy- and deoxyhaemoglobin with high technical reproducibility (< 4% variability) (88, 89). Similarly, mesoscopic PAI enables only the characterisation of superficial (*approx* 4 mm) perfused vasculature *in vivo*. The full extent of the investigated tumour vascular networks – especially in perfusion-limited vessels – is not fully captured in this detection configuration. For instance, due to the top illumination and planar detection geometry, vessels approaching 90° to the detection plane are not detected, creating discontinuities in the network and artificially increasing the number of CC (90). Thus, the quantitative biomarkers presented in this study cannot be taken outside their context or as absolute measurements. Tailored contrast agents that have been introduced recently could be employed for identifying specifically cellular response mechanisms such as senescence (91, 92).

A second key limitation is that we have not completed serial culling of animals to obtain tumour samples at all imaging time-points of interest. Instead, we favoured having paired longitudinal imaging data to improve statistical power, thus decreasing the need for high animal numbers per group. Such *ex vivo* IHC analysis would have provided time-resolved validation of our *in vivo* imaging. Furthermore, we did not keep animals alive longer than 7 days post-RT on average to further monitor tumour growth, vascular alterations, or survival. Thus, considering the time-scale of tumour and vascular response to RT, our imaging-based investigation depicts the earlier phase of response that may not recapitulate the full story, especially in terms of recurrence or revascularisation post-RT (93). Nevertheless, targeted neoadjuvant RT did improve resectability at endpoint given the decrease in tumour size.

Finally, the imaging time-points in terms of days between treated groups were different due to the nature of the hypofractionated (HFRT) vs. single-dose (SDRT) courses of treatment. Even though treatment arms are compared directly, they are only comparable in terms of total dose delivered but not in absolute number of days. Further study would be required to control for the time effect more directly. We believe that changes in intratumoural blood oxygen saturation following RT have an important dose- and time-dependence, which could be tumour-specific, as highlighted in our comparisons with prior literature. We acknowledge that there is a time compartment to radiation response, but report here differences related to the impact of fractionation and dose per fraction.

In terms of clinical translatability, although neoadjuvant RT is not the main line of treatment in breast cancer patients, it has been shown to be beneficial in low-risk or early-stage breast cancer, including for ER+ cases with potential chemoresistance (94–96). Indeed, there has been an increasing interest in breast cancer pre-operative hypofractionated RT or even SDRT, enabled by the increased precision in tumour targeting provided by recent developments in integrated imaging and accelerator technologies (*e*.*g*. magnetic resonance-guided RT, PET-guided – also termed biology-guided – RT, or camera-guided CyberKnife) that decrease overall treatment time (97). Pre-operative partial breast irradiation using 20 Gy SDRT to the tumour volume, similar to that described in our preclinical study, has been investigated and shown oncologically safe in a feasibility study and a 5-year follow-up study (61, 62), and in another trial using 21 Gy SDRT (63), and are both being further investigated in on-going clinical trials (ClinicalTrials.gov ID, NCT03863301, NCT02212860, and NCT04679454; Australian New Zealand Clinical Trials Registry, ACTRN12621000659808) (64, 65). The potential of PAI in this context is multi-faceted, and clinical studies should investigate the use of quantitative PAI in breast cancer patients receiving RT or combination therapies (38). For instance, preclinical data in numerous cancer models already support the benefits of using functional imaging modalities to guide the boosting of the radiation dose to tumour hypoxic fraction, using oxygen-guided spatially modulated dose delivery (14, 98). Thus, the role of label-free portable bedside clinical photoacoustic imagers in the radiation oncology clinics should be further investigated, especially in the context of treatment adaptation.

## Conclusion

We demonstrated that PAI reveals the differential effect between hypofractionated and ablative courses of RT in two breast cancer models, with increased blood content and gas challenge response pre-RT associated with improved treatment outcomes. We showed that PAI could capture early RT response, days before any change in tumour volume was measured, informing on radioresistance. The vascular alterations resulting from ablative doses of radiation were seen within 24h of treatment in the more radiosensitive model. Hence, the molecular contrast provided by PAI shows promise for measuring functional response and shedding new light on the vascular compartment of radiation response, as well as the impact of hypoxia on radioresistance *in vivo*. We envision that PAI biomarkers will benefit other cancer types treated with RT, such as already demonstrated preclinically in head & neck cancer (44, 45), with high translational potential in those more superficial tumour sites. Visualising and quantifying the tumour vascular phenotype *in vivo* thus has the potential to enable personalised RT through biology-tailored targeting and further inform the course of treatment.

## ACKNOWLEDGEMENTS

TLL, MEO, EVB, TRE, LW, MAG, LH, CB, PWS, and SEB acknowledge the support from Cancer Research UK (CRUK) under grant numbers C14303/A17197, C9545/A29580, C47594/A16267, C197/A16465, C47594/A29448, and CRUK RadNet Cambridge under the grant number C17918/A28870. SEB also receives funding support from the UKRI Engineering and Physical Sciences Research Council (EP/R003599/1, EP/X037770/1 and EP/V027069/1). TLL is supported by the Cambridge Trust. EVB is supported by CRUK RadNet. LH acknowledges research support from Against Breast Cancer. We thank the CRUK Cambridge Institute’s Biological Resources Unit, Imaging Core, Histopathological Core, Light Microscopy, Research Instrumentation and Cell Services, and the RadNet Small Animal Hospital for their support in conducting this research. We also acknowledge the CRUK RadNet leadership team, Prof. Charlotte Coles and Nicola Leblond, for infrastructure support. We would like to thank Prof. Raj Jena and Dr. Florian Markowetz for useful discussions and guidance throughout study conduction.

## AUTHOR CONTRIBUTIONS

TLL: conceptualisation, data curation, formal analysis, funding acquisition, investigation, methodology, software, validation, visualisation, writing - original draft. M-EO: investigation, methodology, project administration, supervision, validation, writing - review & editing. EVB: investigation, methodology, validation, writing - review & editing. TRE: methodology, software, validation, writing - review & editing. LW: investigation, validation, writing - review & editing. MAG: investigation, writing - review & editing. LH: investigation, validation, writing - review & editing. CB: formal analysis, investigation, methodology, software, writing - review & editing. SK: investigation, methodology, writing - review & editing. YC: investigation, methodology, writing - review & editing. LY: investigation, methodology, resources, project administration, writing - review & editing. PWS: formal analysis, investigation, methodology, software, supervision, validation, visualisation, writing - review & editing. SEB: conceptualisation, funding acquisition, methodology, project administration, resources, supervision, validation, writing - review & editing.

## COMPETING FINANCIAL INTERESTS

The authors have no conflict of interest related to the present manuscript to disclose.

## Notes

https://doi.org/10.17863/CAM.118247

### Competing Interest Statement

The authors have declared no competing interest.

## Bibliography

1. Mark W. Dewhirst, Yiting Cao, and Benjamin Moeller. Cycling hypoxia and free radicals regulate angiogenesis and radiotherapy response. Nature Reviews Cancer, 8(6):425–437, 2008.

2. Katheryn Begg and Mahvash Tavassoli. Inside the hypoxic tumour: reprogramming of the ddr and radioresistance. Cell Death Discovery, 6(1):77, 2020.

3. M. R. Horsman and J. Overgaard. The impact of hypoxia and its modification of the outcome of radiotherapy. J Radiat Res, 57 Suppl 1(Suppl 1):i90–i98, 2016.

4. G. L. Semenza. The hypoxic tumor microenvironment: A driving force for breast cancer progression. Biochim Biophys Acta, 1863(3):382–391, 2016.

5. Claudia Sousa, Mafalda Cruz, Ana Neto, Kayla Pereira, Marta Peixoto, Joana Bastos, Monica Henriques, Domingos Roda, Rui Marques, Cristina Miranda, Gilberto Melo, Gabriela Sousa, Paulo Figueiredo, and Paula Alves. Neoadjuvant radiotherapy in the approach of locally advanced breast cancer. ESMO Open, 5(2):e000640, 2020.

6. Mathilde Maire, Marc Debled, Adeline Petit, Marion Fournier, Gaetan Macgrogan, Nathalie Quenel-Thueux, Helene Charitansky, Simone Mathoulin-Pelissier, Herve Bonnefoi, and Christine Tunon de Lara. Neoadjuvant chemotherapy and radiotherapy for locally advanced breast cancer: Safety and efficacy of reverse sequence compared to standard technique? European Journal of Surgical Oncology, 48(8):1699–1705, 2022.

7. Phoebe Chidley, Farshad Foroudi, Mark Tacey, Richard Khor, Janice Yeh, Elaine Bevington, Anthony Hyett, Su Wen Loh, Grace Chew, James McCracken, Derek Neoh, Belinda Yeo, Caroline Baker, Sunil Jassal, Michael Law, Natalie Zantuck, Margaret Cokelek, Mario Guerrieri, Belinda Brown, David Stoney, Michael Ng, and Michael Chao. Neoadjuvant radiotherapy for locally advanced and high-risk breast cancer. Journal of Medical Imaging and Radiation Oncology, 65(3):345–353, 2021.

8. Yuan-Hong Lin, Phoebe Chidley, Lorenztino Admojo, Sunil Jassal, Natalie Zantuck, Farshad Foroudi, Elaine Bevington, Grace Chew, Anthony Hyett, Su Wen Loh, Suat Li Ng, Tristan Leech, Caroline Baker, Michael Law, Wei Ming Ooi, Charles Yong, Richard Khor, and Michael Chao. Pathologic complete response and oncologic outcomes in locally advanced breast cancers treated with neoadjuvant radiation therapy: An australian perspective. Practical Radiation Oncology, 13(4):301–313, 2023.

9. R. H. Thomlinson and L. H. Gray. The histological structure of some human lung cancers and the possible implications for radiotherapy. British Journal of Cancer, 9(4):539–549, 1955.

10. D. R. Grimes and M. Partridge. A mechanistic investigation of the oxygen fixation hypothesis and oxygen enhancement ratio. Biomed Phys Eng Express, 1(4):045209, 2015.

11. R. A. Chandra, F. K. Keane, F. E. M. Voncken, and Jr. Thomas, C.R. Contemporary radiotherapy: present and future. Lancet, 398(10295):171–184, 2021.

12. J. M. Brown, M. Diehn, and Jr. Loo, B.W. Stereotactic ablative radiotherapy should be combined with a hypoxic cell radiosensitizer. Int J Radiat Oncol Biol Phys, 78(2):323–7, 2010.

13. J. M. Brown, D. J. Carlson, and D. J. Brenner. The tumor radiobiology of srs and sbrt: are more than the 5 rs involved? Int J Radiat Oncol Biol Phys, 88(2):254–62, 2014.

14. B. Epel, M. C. Maggio, E. D. Barth, R. C. Miller, C. A. Pelizzari, M. Krzykawska-Serda, S. V. Sundramoorthy, B. Aydogan, R. R. Weichselbaum, V. M. Tormyshev, and H. J. Halpern. Oxygen-guided radiation therapy. Int J Radiat Oncol Biol Phys, 103(4):977–984, 2019.

15. Z. Fuks and R. Kolesnick. Engaging the vascular component of the tumor response. Cancer Cell, 8(2):89–91, 2005.

16. H. J. Park, R. J. Griffin, S. Hui, S. H. Levitt, and C. W. Song. Radiation-induced vascular damage in tumors: implications of vascular damage in ablative hypofractionated radiotherapy (sbrt and srs). Radiat Res, 177(3):311–27, 2012.

17. C. W. Song, Y. J. Lee, R. J. Griffin, I. Park, N. A. Koonce, S. Hui, M. S. Kim, K. E. Dusenbery, P. W. Sperduto, and L. C. Cho. Indirect tumor cell death after high-dose hypofractionated irradiation: Implications for stereotactic body radiation therapy and stereotactic radiation surgery. Int J Radiat Oncol Biol Phys, 93(1), 2015.

18. D. Klein. The tumor vascular endothelium as decision maker in cancer therapy. Front Oncol, 8:367, 2018.

19. R. Kolesnick and Z. Fuks. Radiation and ceramide-induced apoptosis. Oncogene, 22(37):5897–906, 2003.

20. M. Garcia-Barros, F. Paris, C. Cordon-Cardo, D. Lyden, S. Rafii, A. Haimovitz-Friedman, Z. Fuks, and R. Kolesnick. Tumor response to radiotherapy regulated by endothelial cell apoptosis. Science, 300(5622):1155–9, 2003.

21. R. Jared Weinfurtner, Natarajan Raghunand, Olya Stringfield, Mahmoud Abdalah, Bethany L. Niell, Dana Ataya, Angela Williams, Blaise Mooney, Marilin Rosa, Marie C. Lee, Nazanin Khakpour, Christine Laronga, Brian Czerniecki, Roberto Diaz, Kamran Ahmed, Iman Washington, and Michael Montejo. Mri response to pre-operative stereotactic ablative body radiotherapy (sabr) in early stage er/pr+ her2-breast cancer correlates with surgical pathology tumor bed cellularity. Clinical Breast Cancer, 22(2):e214–e223, 2022.

22. Ayyaz Qadir, Nabita Singh, Aung Aung Kywe Moe, Glenn Cahoon, Jessica Lye, Michael Chao, Farshad Foroudi, and Sergio Uribe. Potential of mri in assessing treatment response after neoadjuvant radiation therapy treatment in breast cancer patients: A scoping review. Clinical Breast Cancer, 2024.

23. D. R. Grimes, D. R. Warren, and S. Warren. Hypoxia imaging and radiotherapy: bridging the resolution gap. Br J Radiol, 90(1076):20160939, 2017.

24. R. A. D’Alonzo, S. Gill, P. Rowshanfarzad, S. Keam, K. M. MacKinnon, A. M. Cook, and M. A. Ebert. In vivo noninvasive preclinical tumor hypoxia imaging methods: a review. Int J Radiat Biol, 97 (5):593–631, 2021.

25. I. Toma-Dasu, J. Uhrdin, L. Antonovic, A. Dasu, S. Nuyts, P. Dirix, K. Haustermans, and A. Brahme. Dose prescription and treatment planning based on fmiso-pet hypoxia. Acta Oncol, 51(2):222–30, 2012.

26. N. Y. Lee, J. G. Mechalakos, S. Nehmeh, Z. Lin, O. D. Squire, S. Cai, K. Chan, P. B. Zanzonico, C. Greco, C. C. Ling, J. L. Humm, and H. Schoder. Fluorine-18-labeled fluoromisonidazole positron emission and computed tomography-guided intensity-modulated radiotherapy for head and neck cancer: a feasibility study. Int J Radiat Oncol Biol Phys, 70(1):2–13, 2008.

27. N. Lee, H. Schoder, B. Beattie, R. Lanning, N. Riaz, S. McBride, N. Katabi, D. Li, B. Yarusi, S. Chan, L. Mitrani, Z. Zhang, D. G. Pfister, E. Sherman, S. Baxi, J. Boyle, L. G. Morris, I. Ganly, R. Wong, and J. Humm. Strategy of using intratreatment hypoxia imaging to selectively and safely guide radiation dose de-escalation concurrent with chemotherapy for locoregionally advanced human papillomavirus-related oropharyngeal carcinoma. Int J Radiat Oncol Biol Phys, 96(1):9–17, 2016.

28. Rami R. Hallac, Heling Zhou, Rajesh Pidikiti, Kwang Song, Strahinja Stojadinovic, Dawen Zhao, Timothy Solberg, Peter Peschke, and Ralph P. Mason. Correlations of noninvasive bold and told mri with po2 and relevance to tumor radiation response. Magnetic Resonance in Medicine, 71(5):1863–1873, 2014.

29. Andreas Stadlbauer, Max Zimmermann, Barbara Bennani-Baiti, Thomas H Helbich, Pascal Baltzer, Paola Clauser, Panagiotis Kapetas, Zsuzsanna Bago-Horvath, and Katja Pinker. Development of a non-invasive assessment of hypoxia and neovascularization with magnetic resonance imaging in benign and malignant breast tumors: initial results. Molecular Imaging and Biology, 21:758–770, 2019.

30. N. Wiedenmann, A. L. Grosu, M. Buchert, H. C. Rischke, J. Ruf, L. Bielak, L. Majerus, A. Ruhle, F. Bamberg, D. Baltas, J. Hennig, M. Mix, M. Bock, and N. H. Nicolay. The utility of multiparametric mri to characterize hypoxic tumor subvolumes in comparison to fmiso pet/ct. consequences for diagnosis and chemoradiation treatment planning in head and neck cancer. Radiother Oncol, 150:128–135, 2020.

31. T. Hompland, K. H. Hole, H. B. Ragnum, E. K. Aarnes, L. Vlatkovic, A. K. Lie, S. Patzke, B. Brennhovd, T. Seierstad, and H. Lyng. Combined mr imaging of oxygen consumption and supply reveals tumor hypoxia and aggressiveness in prostate cancer patients. Cancer Res, 78(16):4774–4785, 2018.

32. T. Hillestad, T. Hompland, C. S. Fjeldbo, V. E. Skingen, U. B. Salberg, E. K. Aarnes, A. Nilsen, K. V. Lund, T. S. Evensen, G. B. Kristensen, T. Stokke, and H. Lyng. Mri distinguishes tumor hypoxia levels of different prognostic and biological significance in cervical cancer. Cancer Res, 80(18):3993–4003, 2020.

33. Anna Li, Erlend Andersen, Christoffer Lervåg, Cathinka H. Julin, Heidi Lyng, Taran P. Hellebust, and Eirik Malinen. Dynamic contrast enhanced magnetic resonance imaging for hypoxia mapping and potential for brachytherapy targeting. Physics and Imaging in Radiation Oncology, 2:1–6, 2017.

34. J. P. Heiken. Contrast safety in the cancer patient: preventing contrast-induced nephropathy. Cancer Imaging, 8 Spec No A(Spec Iss A):S124–7, 2008.

35. M. Rogosnitzky and S. Branch. Gadolinium-based contrast agent toxicity: a review of known and proposed mechanisms. Biometals, 29(3):365–76, 2016.

36. C. Michiels, C. Tellier, and O. Feron. Cycling hypoxia: A key feature of the tumor microenvironment. Biochim Biophys Acta, 1866(1):76–86, 2016.

37. P. N. Span and J. Bussink. Biology of hypoxia. Semin Nucl Med, 45(2):101–9, 2015.

38. T. L. Lefebvre, E. Brown, L. Hacker, T. Else, M. E. Oraiopoulou, M. R. Tomaszewski, R. Jena, and S. E. Bohndiek. The potential of photoacoustic imaging in radiation oncology. Front Oncol, 12:803777, 2022.

39. M. Omar, J. Aguirre, and V. Ntziachristos. Optoacoustic mesoscopy for biomedicine. Nat Biomed Eng, 3(5):354–370, 2019.

40. Michaela Taylor-Williams, Graham Spicer, Gemma Bale, and Sarah E. Bohndiek. Noninvasive hemoglobin sensing and imaging: optical tools for disease diagnosis. Journal of Biomedical Optics, 27(8):080901, 2022.

41. E. Brown, J. Brunker, and S. E. Bohndiek. Photoacoustic imaging as a tool to probe the tumour microenvironment. Dis Model Mech, 12(7), 2019.

42. M. R. Tomaszewski, I. Q. Gonzalez, J. P. O’Connor, O. Abeyakoon, G. J. Parker, K. J. Williams, F. J. Gilbert, and S. E. Bohndiek. Oxygen enhanced optoacoustic tomography (oe-ot) reveals vascular dynamics in murine models of prostate cancer. Theranostics, 7(11):2900–2913, 2017.

43. M. Omar, J. Gateau, and V. Ntziachristos. Raster-scan optoacoustic mesoscopy in the 25-125 mhz range. Opt Lett, 38(14):2472–4, 2013.

44. L. J. Rich and M. Seshadri. Photoacoustic monitoring of tumor and normal tissue response to radiation. Sci Rep, 6:21237, 2016.

45. L. J. Rich, A. Miller, A. K. Singh, and M. Seshadri. Photoacoustic imaging as an early biomarker of radio therapeutic efficacy in head and neck cancer. Theranostics, 8(8):2064–2078, 2018.

46. Anna Orlova, Ksenia Pavlova, Aleksey Kurnikov, Anna Maslennikova, Marina Myagcheva, Evgeniy Zakharov, Dmitry Skamnitskiy, Valeria Perekatova, Alexander Khilov, Andrey Kovalchuk, Alexander Moiseev, Ilya Turchin, Daniel Razansky, and Pavel Subochev. Noninvasive optoacoustic microangiography reveals dose and size dependency of radiation-induced deep tumor vasculature remodeling. Neoplasia, 26:100778, 2022.

47. Zhongli Cai, Zhuo Chen, Kristy E. Bailey, Deborah A. Scollard, Raymond M. Reilly, and Katherine A. Vallis. Relationship between induction of phosphorylated h2ax and survival in breast cancer cells exposed to 111in-dtpa-hegf. Journal of Nuclear Medicine, 49(8):1353–1361, 2008.

48. Steven Eschrich, Hongling Zhang, Haiyan Zhao, David Boulware, Ji-Hyun Lee, Gregory Bloom, and Javier F. Torres-Roca. Systems biology modeling of the radiation sensitivity network: A biomarker discovery platform. International Journal of Radiation Oncology*Biology*Physics, 75(2):497–505, 2009.

49. M. Villalobos. Radiosensitivity of human breast cancer cell lines of different hormonal responsiveness. modulatory effects of oestradiol. International Journal of Radiation Biology, 70(2):161–169, 1996.

50. George P. Amorino, Michael L. Freeman, and Hak Choy. Enhancement of Radiation Effects In Vitro by the Estrogen Metabolite 2-Methoxyestradiol. Radiation Research, 153(4):384 – 391, 2000.

51. I. Quiros-Gonzalez, M. R. Tomaszewski, S. J. Aitken, L. Ansel-Bollepalli, L. A. McDuffus, M. Gill, L. Hacker, J. Brunker, and S. E. Bohndiek. Optoacoustics delineates murine breast cancer models displaying angiogenesis and vascular mimicry. Br J Cancer, 118(8):1098–1106, 2018.

52. P. Rinwa, M. Eriksson, I. Cotgreave, and M. Bäckberg. 3r-refinement principles: elevating rodent well-being and research quality. Lab Anim Res, 40(1):11, 2024.

53. Sivanandane Sittadjody. Controlled release of hormones by pellet implants. In Controlled Drug Delivery Systems, pages 91–107. CRC Press, 2020.

54. K. V. Gates, E. Alamaw, K. Jampachaisri, M. K. Huss, and C. Pacharinsak. Efficacy of supplemental diet gels for preventing postoperative weight loss in mice (mus musculus). J Am Assoc Lab Anim Sci, 62(1):87–91, 2023.

55. C. M. Ma, C. W. Coffey, L. A. DeWerd, C. Liu, R. Nath, S. M. Seltzer, J. P. Seuntjens, and Medicine American Association of Physicists in. Aapm protocol for 40-300 kv x-ray beam dosimetry in radiotherapy and radiobiology. Med Phys, 28(6):868–93, 2001.

56. B. Fraass, K. Doppke, M. Hunt, G. Kutcher, G. Starkschall, R. Stern, and J. Van Dyke. American association of physicists in medicine radiation therapy committee task group 53:quality assurance for clinical radiotherapy treatment planning. Med Phys, 25(10):1773–829, 1998.

57. P. E. Lindsay, P. V. Granton, A. Gasparini, S. Jelveh, R. Clarkson, S. van Hoof, J. Hermans, J. Kaas, F. Wittkamper, J. J. Sonke, F. Verhaegen, and D. A. Jaffray. Multi-institutional dosimetric and geometric commissioning of image-guided small animal irradiators. Med Phys, 41(3):031714, 2014.

58. J. Wong, E. Armour, P. Kazanzides, I. Iordachita, E. Tryggestad, H. Deng, M. Matinfar, C. Kennedy, Z. Liu, T. Chan, O. Gray, F. Verhaegen, T. McNutt, E. Ford, and T. L. DeWeese. High-resolution, small animal radiation research platform with x-ray tomographic guidance capabilities. Int J Radiat Oncol Biol Phys, 71(5):1591–9, 2008.

59. Robert Jacques, Russell Taylor, John Wong, and Todd McNutt. Towards real-time radiation therapy: Gpu accelerated superposition/convolution. Computer Methods and Programs in Biomedicine, 98(3):285–292, 2010.

60. Robert Jacques, John Wong, Russell Taylor, and Todd McNutt. Real-time dose computation: Gpu-accelerated source modeling and superposition/convolution. Medical Physics, 38(1):294–305, 2011.

61. Danny Lavigne, Tarek Hijal, Peter Vavassis, Marie-Christine Guilbert, Lucas Sideris, Pierre Dubé, Mai-Kim Gervais, Guy Leblanc, Michel-Pierre Dufresne, David Nguyen, David Tiberi, Dima Mahmoud, and Michael Yassa. Single preoperative radiation therapy with delayed surgery for low-risk breast cancer: Oncologic outcome, toxicity and cosmesis of the sport-ds phase i trial. Radiotherapy and Oncology, 200:110515, 2024.

62. Yasmin A. Civil, Jeanine E. Vasmel, Ramona K. Charaghvandi, Anette C. Houweling, Celien P.H. Vreuls, Paul J. van Diest, Arjen J. Witkamp, Annemiek Doeksen, Thijs van Dalen, Joeke Felderhof, Iris van Dam, Ben J. Slotman, Anna M. Kirby, Helena M. Verkooijen, Susanne van der Velde, Femke van der Leij, and H.J.G. Desiree van den Bongard. Preoperative magnetic resonance guided single-dose partial breast irradiation: 5-year results of the prospective single-arm ablative trial. International Journal of Radiation Oncology*Biology*Physics, 2024.

63. K. Guidolin, B. Yaremko, K. Lynn, S. Gaede, A. Kornecki, G. Muscedere, I. BenNachum, O. Shmuilovich, M. Mouawad, E. Yu, T. Sexton, N. Gelman, V. Moiseenko, M. Brackstone, and M. Lock. Stereotactic image-guided neoadjuvant ablative single-dose radiation, then lumpectomy, for early breast cancer: the signal prospective single-arm trial of single-dose radiation therapy. Curr Oncol, 26(3):e334–e340, 2019.

64. Yasmin A. Civil, Arlene L. Oei, Katya M. Duvivier, Nina Bijker, Philip Meijnen, Lorraine Donkers, Sonja Verheijen, Zdenko van Kesteren, Miguel A. Palacios, Laura J. Schijf, Ellis Barba, Inge R. H. M. Konings, C. Willemien Menke van der Houven van Oordt, Paulien G. Westhoff, Hanneke J. M. Meijer, Gwen M. P. Diepenhorst, Victor Thijssen, Florent Mouliere, Berend J. Slotman, Susanne van der Velde, and H. J. G. Desiree van den Bongard. Prediction of pathologic complete response after single-dose mr-guided partial breast irradiation in low-risk breast cancer patients: the ablative-2 trial: a study protocol. BMC Cancer, 23(1):419, 2023.

65. Maria Alessia Zerella, Mattia Zaffaroni, Giuseppe Ronci, Samantha Dicuonzo, Damaris Patricia Rojas, Anna Morra, Cristiana Fodor, Elena Rondi, Sabrina Vigorito, Francesca Botta, Marta Cremonesi, Cristina Garibaldi, Silvia Penco, Viviana Enrica Galimberti, Mattia Intra, Sara Gandini, Massimo Barberis, Giuseppe Renne, Federica Cattani, Paolo Veronesi, Roberto Orecchia, Barbara Alicja Jereczek-Fossa, and Maria Cristina Leonardi. Single fraction ablative preoperative radiation treatment for early-stage breast cancer: the crystal study a phase i/ii clinical trial protocol. BMC Cancer, 22(1):358, 2022.

66. A. Murray Brunt, J. S. Haviland, D. A. Wheatley, M. A. Sydenham, A. Alhasso, D. J. Bloomfield, C. Chan, M. Churn, S. Cleator, C. E. Coles, A. Goodman, A. Harnett, P. Hopwood, A. M. Kirby, C. C. Kirwan, C. Morris, Z. Nabi, E. Sawyer, N. Somaiah, L. Stones, I. Syndikus, J. M. Bliss, J. R. Yarnold, and F. AST-Forward Trial Management Group. Hypofractionated breast radiotherapy for 1 week versus 3 weeks (fast-forward): 5-year efficacy and late normal tissue effects results from a multicentre, non-inferiority, randomised, phase 3 trial. Lancet, 395(10237):1613–1626, 2020.

67. S. Park, A. B. Karpiouk, S. R. Aglyamov, and S. Y. Emelianov. Adaptive beamforming for photoacoustic imaging. Opt Lett, 33(12):1291–3, 2008.

68. M. Omar, D. Soliman, J. Gateau, and V. Ntziachristos. Ultrawideband reflection-mode optoacoustic mesoscopy. Opt Lett, 39(13):3911–4, 2014.

69. Paul W. Sweeney, Lina Hacker, Thierry L. Lefebvre, Emma L. Brown, Janek Grohl, and Sarah E. Bohndiek. Unsupervised segmentation of 3d microvascular photoacoustic images using deep generative learning. Advanced Science, 11(32):2402195, 2024.

70. E. L. Brown, T. L. Lefebvre, P. W. Sweeney, B. J. Stolz, J. Grohl, L. Hacker, Z. Huang, D. L. Couturier, H. A. Harrington, H. M. Byrne, and S. E. Bohndiek. Quantification of vascular networks in photoacoustic mesoscopy. Photoacoustics, 26:100357, 2022.

71. Bernadette J. Stolz, Jakob Kaeppler, Bostjan Markelc, Franziska Braun, Florian Lipsmeier, Ruth J. Muschel, Helen M. Byrne, and Heather A. Harrington. Multiscale topology characterizes dynamic tumor vascular networks. Science Advances, 8(23):eabm2456, 2022.

72. Thomas R. Else, Janek Grohl, Lina Hacker, and Sarah E. Bohndiek. Patato: a python photoacoustic tomography analysis toolkit. Journal of Open Source Software, 9(93):5686, 2024.

73. Sture Holm. A simple sequentially rejective multiple test procedure. Scandinavian Journal of Statistics, 6(2):65–70, 1979.

74. E. Draeger, A. Sawant, C. Johnstone, B. Koger, S. Becker, Z. Vujaskovic, I. L. Jackson, and Y. Poirier. A dose of reality: How 20 years of incomplete physics and dosimetry reporting in radiobiology studies may have contributed to the reproducibility crisis. Int J Radiat Oncol Biol Phys, 106(2):243–252, 2020.

75. Silvester J. Bartsch, Klara Brozova, Viktoria Ehret, Joachim Friske, Christoph Furbock, Lukas Kenner, Daniela Laimer-Gruber, Thomas H. Helbich, and Katja Pinker. Non-contrast-enhanced multiparametric mri of the hypoxic tumor microenvironment allows molecular subtyping of breast cancer: A pilot study. Cancers, 16(2), 2024.

76. Chen-Ting Lee, Yingchun Zhou, Kingshuk Roy-Choudhury, Sharareh Siamakpour-Reihani, Kenneth Young, Peter Hoang, John P. Kirkpatrick, Jen-Tsan A. Chi, Mark W. Dewhirst, and Janet K. Horton. Subtype-Specific Radiation Response and Therapeutic Effect of FAS Death Receptor Modulation in Human Breast Cancer. Radiation Research, 188(2):169 – 180, 2017.

77. J. M. Fleming, T. C. Miller, M. J. Meyer, E. Ginsburg, and B. K. Vonderhaar. Local regulation of human breast xenograft models. J Cell Physiol, 224(3):795–806, 2010.

78. Márcia Martinho Costa, Anant Shah, Ian Rivens, Carol Box, Tuathan O’Shea, Jeffrey Bamber, and Gail ter Haar. Photoacoustic imaging for the prediction and assessment of response to radiotherapy in vivo. bioRxiv, page 329516, 2018.

79. Shreya Goel, Jorge de la Cerda, William Schuler, Aikaterini Kotrotsou, Julio Cardenas-Rodriguez, and Mark Pagel. Evaluations of radiotherapy in small animal models of pancreatic cancer with oxygen enhanced–dynamic contrast enhanced multispectral optoacoustic tomography (OE-DCE MSOT). In Photons Plus Ultrasound: Imaging and Sensing 2023, volume 12379, page 123791B. International Society for Optics and Photonics, SPIE, 2023. doi: 10.1117/12.2648074.

80. Janggun Jo, Jeff Folz, Maria E. Gonzalez, Alessandro Paolo, Ahmad Eido, Eamon Salfi, Shilpa Tekula, Sebastiano And, Roberta Caruso, Celina G. Kleer, Xueding Wang, and Raoul Kopelman. Personalized oncology by in vivo chemical imaging: Photoacoustic mapping of tumor oxygen predicts radiotherapy efficacy. ACS Nano, 17(5):4396–4403, 2023.

81. S. Goel, J. de la Cerda, W. Schuler, A. Kotrotsou, J. Cárdenas-Rodríguez, and M. D. Pagel. Improving evaluations of radiation therapy with dynamic contrast enhanced multispectral optoacoustic tomography (dce msot). Proceedings of the European Molecular Imaging Meeting, pages PS 09–03, 2020.

82. J. Bussink, J. H A. M. Kaanders, P. F J. W. Rijken, J. A. Raleigh, and A. J. Van der Kogel. Changes in blood perfusion and hypoxia after irradiation of a human squamous cell carcinoma xenograft tumor line. Radiation Research, 153(4):398–404, 2000.

83. J. Kory, V. Narain, B. J. Stolz, J. Kaeppler, B. Markelc, R. J. Muschel, P. K. Maini, J. M. Pitt-Francis, and H. M. Byrne. Enhanced perfusion following exposure to radiotherapy: A theoretical investigation. PLoS Comput Biol, 20(2):e1011252, 2024.

84. A. Orlova, M. Sirotkina, E. Smolina, V. Elagin, A. Kovalchuk, I. Turchin, and P. Subochev. Raster-scan optoacoustic angiography of blood vessel development in colon cancer models. Photoacoustics, 13:25–32, 2019.

85. Jakob R Kaeppler, Jianzhou Chen, Mario Buono, Jenny Vermeer, Pavitra Kannan, Weiâ CChen Cheng, Dimitrios Voukantsis, James M Thompson, Mark A Hill, Danny Allen, Ana Gomes, Veerle Kersemans, Paul Kinchesh, Sean Smart, Francesca Buffa, Claus Nerlov, Ruth J Muschel, and Bostjan Markelc. Endothelial cell death after ionizing radiation does not impair vascular structure in mouse tumor models. EMBO reports, 23(9):e53221, 2022.

86. E. Hysi, M. N. Fadhel, Y. Wang, J. A. Sebastian, A. Giles, G. J. Czarnota, A. A. Exner, and M. C. Kolios. Photoacoustic imaging biomarkers for monitoring biophysical changes during nanobubble-mediated radiation treatment. Photoacoustics, 20:100201, 2020.

87. Xu Cao, Srinivasa Rao Allu, Shudong Jiang, Mengyu Jia, Jason R. Gunn, Cuiping Yao, Ethan P. LaRochelle, Jennifer R. Shell, Petr Bruza, David J. Gladstone, Lesley A. Jarvis, Jie Tian, Sergei A. Vinogradov, and Brian W. Pogue. Tissue po2 distributions in xenograft tumors dynamically imaged by cherenkov-excited phosphorescence during fractionated radiation therapy. Nature Communications, 11(1):573, 2020.

88. J. Joseph, M. R. Tomaszewski, I. Quiros-Gonzalez, J. Weber, J. Brunker, and S. E. Bohndiek. Evaluation of precision in optoacoustic tomography for preclinical imaging in living subjects. J Nucl Med, 58(5):807–814, 2017.

89. Y. Poirier, C. D. Johnstone, A. Anvari, N. P. Brodin, M. D. Santos, M. Bazalova-Carter, and A. Sawant. A failure modes and effects analysis quality management framework for image-guided small animal irradiators: A change in paradigm for radiation biology. Med Phys, 47(4):2013–2022, 2020.

90. L. Hacker, E. L. Brown, T. L. Lefebvre, P. W. Sweeney, and S. E. Bohndiek. Performance evaluation of mesoscopic photoacoustic imaging. Photoacoustics, 31:100505, 2023.

91. Andrew G. Baker, Muhamad Hartono, Hui-Ling Ou, Andrea BistroviÄ‡ Popov, Emma L. Brown, James Joseph, Monika Golinska, Estela GonzÃ¡lez-Gualda, David Macias, Jianfeng Ge, Mary Denholm, Samir Morsli, Chandan Sanghera, Thomas R. Else, Heather F. Greer, Aude Vernet, Sarah E. Bohndiek, Daniel MuÃ±oz-EspÃn, and Ljiljana Fruk. An indocyanine green-based nanoprobe for in vivo detection of cellular senescence. Angewandte Chemie International Edition, 63(25):e202404885, 2024.

92. M. Hartono, A. G. Baker, T. R. Else, A. S. Evtushenko, S. E. Bohndiek, D. Munoz-Espin, and L. Fruk. Photoacoustic polydopamine-indocyanine green (pda-icg) nanoprobe for detection of senescent cells. Sci Rep, 14(1):29506, 2024.

93. S. V. Kozin, D. G. Duda, L. L. Munn, and R. K. Jain. Neovascularization after irradiation: what is the source of newly formed vessels in recurring tumors? J Natl Cancer Inst, 104(12):899–905, 2012.

94. Jan Poleszczuk, Kimberly Luddy, Lu Chen, Jae K. Lee, Louis B. Harrison, Brian J. Czerniecki, Hatem Soliman, and Heiko Enderling. Neoadjuvant radiotherapy of early-stage breast cancer and long-term disease-free survival. Breast Cancer Research, 19(1):75, 2017.

95. M. Ahmed, F. Jozsa, and M. Douek. A systematic review of neo-adjuvant radiotherapy in the treatment of breast cancer. Ecancermedicalscience, 15:1175, 2021.

96. Yasmin A. Civil, Lysanne W. Jonker, Maartje P. M. Groot Koerkamp, Katya M. Duvivier, Ralph de Vries, Arlene L. Oei, Berend J. Slotman, Susanne van der Velde, and H. J. G. Desiree van den Bongard. Preoperative partial breast irradiation in patients with low-risk breast cancer: A systematic review of literature. Annals of Surgical Oncology, 30(6):3263–3279, 2023.

97. Mateusz Bilski, Katarzyna Konat-Bška, Maria Alessia Zerella, Stefanie Corradini, Marcin Hetna, Maria Cristina Leonardi, Martyna Gruba, Aleksandra Grzywacz, Patrycja Hatala, Barbara Alicja Jereczek-Fossa, Jacek Fijuth, and žukasz Kuncman. Advances in breast cancer treatment: a systematic review of preoperative stereotactic body radiotherapy (sbrt) for breast cancer. Radiation Oncology, 19(1):103, 2024.

98. Inna Gertsenshteyn, Boris Epel, Mihai Giurcanu, Eugene Barth, John Lukens, Kayla Hall, Jenipher Flores Martinez, Mellissa Grana, Matthew Maggio, Richard C. Miller, Subramanian V. Sundramoorthy, Martyna Krzykawska-Serda, Erik Pearson, Bulent Aydogan, Ralph R. Weichselbaum, Victor M. Tormyshev, Mrignayani Kotecha, and Howard J. Halpern. Absolute oxygen-guided radiation therapy improves tumor control in three preclinical tumor models. Frontiers in Medicine, 10, 2023.

