## Supplementary Materials for "Early Radiation Therapy Response Assessment using Multi-scale Photoacoustic Imaging"

### Supplementary Note 1: Enabling clinically relevant preclinical image-guided radiation therapy through dosimetric quality assurance

Absolute dosimetry in reference conditions (Supplementary Figure 1) showed that the dose output factor of the SARRP was within 3.0% of the reference dose at the SARRP's commissioning across three measurement time-points over the years of the study conduction (min-max measured dose error, 1.2-3.0 %; Supplementary Table 1).
Comparing fixed vs. moving arc delivery, the resulting isodose lines with moving arc delivery spanning 135°were more conformal to the tumour and less dose was delivered to surrounding organs at risk (OAR) (Supplementary Figure 2). The 80% isodose line systematically went deeper within healthy organs with the fixed beam delivery, further illustrated in the dose-volume histograms (DVH, Supplementary Figure 3). Fixed beam and moving arc deliveries achieved similar dose distribution, with around 80% of the segmented tumour receiving 100% of the dose, either 5.0 Gy in a one fraction delivery of the HFRT scheme or 20.0 Gy in the SDRT scheme. However, for OARs sparing, 7-8% more of the OAR volume received 50% of the dose with fixed beam than with moving arc delivery, equating to 2.5 Gy for HFRT or 10.0 Gy for SDRT (48% or 57% of OAR volume with fixed beam vs. 23% or 31% of OAR volume with moving arc, respectively; see arrows in Supplementary Figure 3). More alarming was that 21-23% more of OAR volume received 75% of the planned dose when using fixed beam, that is 3.75 or 15.0 Gy (44% or 54% of OAR volume with fixed beam vs. 40% or 50% of OAR volume with moving arc, respectively; see arrows in Supplementary Figure 3). Dose to the skin was always higher than the planned dose (105%, Supplementary Figure 2), which is likely a direct result of the strong backscattering component of 220 kVp beams (Supplementary Table 2). Electrons produced by photoelectric interaction in the tissue are mostly backward-directed and deposit the dose at shallow depths (mean $e^-$ free path = 1-2 mm). Better dose homogeneity was achieved and the higher skin dose was more conformed to the tumour skin rather than to surrounding OAR skin in moving arc plans. Thus, moving arcs were employed for all subsequent *in vivo* RT experiments performed in this study, with the confidence that the delivered dose was within reasonable tolerance threshold (<5%).

**Supplementary Table 1. Absolute dosimetry of the Small Animal Radiation Research Platform.** Raw charge measurements, corrected charge measurements and resulting dose compared to calibration.

| | Repeat 1 | Repeat 2 | Repeat 3 | Reference |
|---|---|---|---|---|
| **Date** | 19/11/2020 | 27/01/2021 | 19/02/2025 | |
| **Exposure time (s) [Repeats]** | 60 [18] | 60 [12] | 60 [12] | |
| **Raw collected charges (nC)** | 64.61 ± 0.03 | 57.59 ± 0.012 | 56.64 ± 0.03 | |
| **Measured correction factors** | 1.002 ± 0.002 | 1.031 ± 0.002 | 0.992 ± 0.003 | |
| **Corrected collected charges (nC)** | 64.74 | 59.94 | 56.06 | |
| **Dose output (Gy)** | 3.36 | 3.09 | 2.92 | 3.28 |
| **Error (%)** | 1.2 | 3.0 | 5.9 | |

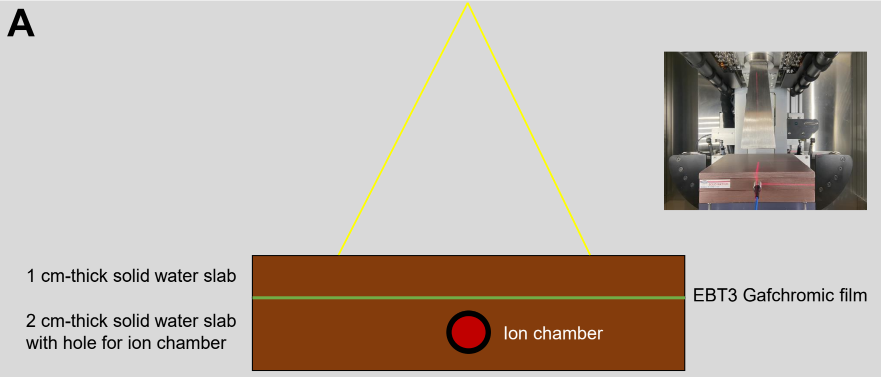


**Supplementary Figure 1. Experimental setup for the dosimetric survey of the SARRP.** A) Dosimetry in reference standard conditions. Solid water phantoms are positioned with fixed source-to-axis distance, radiation beam is turned on for a given amount of time, and dose is measured with an ionisation chamber within the phantom insert by collecting charges which amount is corrected by assessing standard correction factors in different irradiation setups.

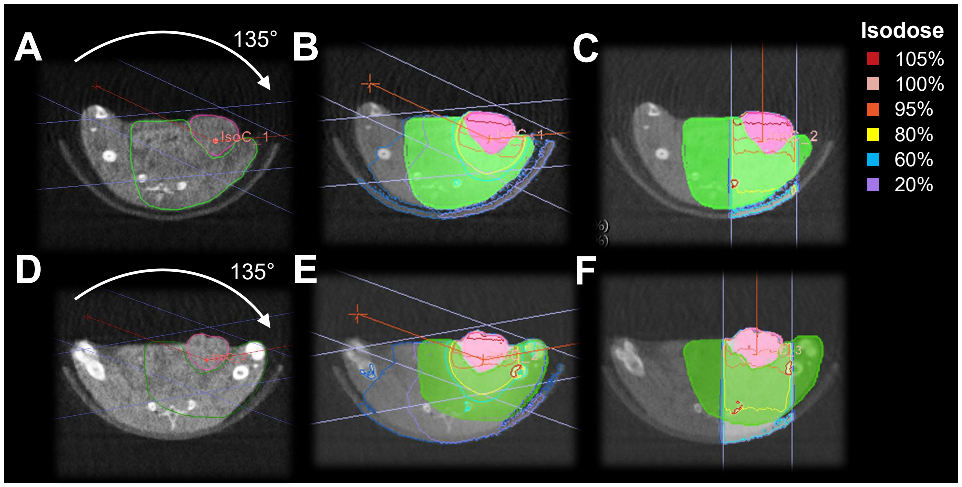


**Supplementary Figure 2. Treatment planning on CBCT axial slices** in A,D) two mice treated with moving arcs. Dose distributions with isodose lines overlaid on CBCT and filled segmentations of the PTV (pink) and of surrounding OAR (green) for two treatment scenarios with the B,E) moving arcs and C,F) fixed beam. Isodose lines represent percent of planned dose to PTV.

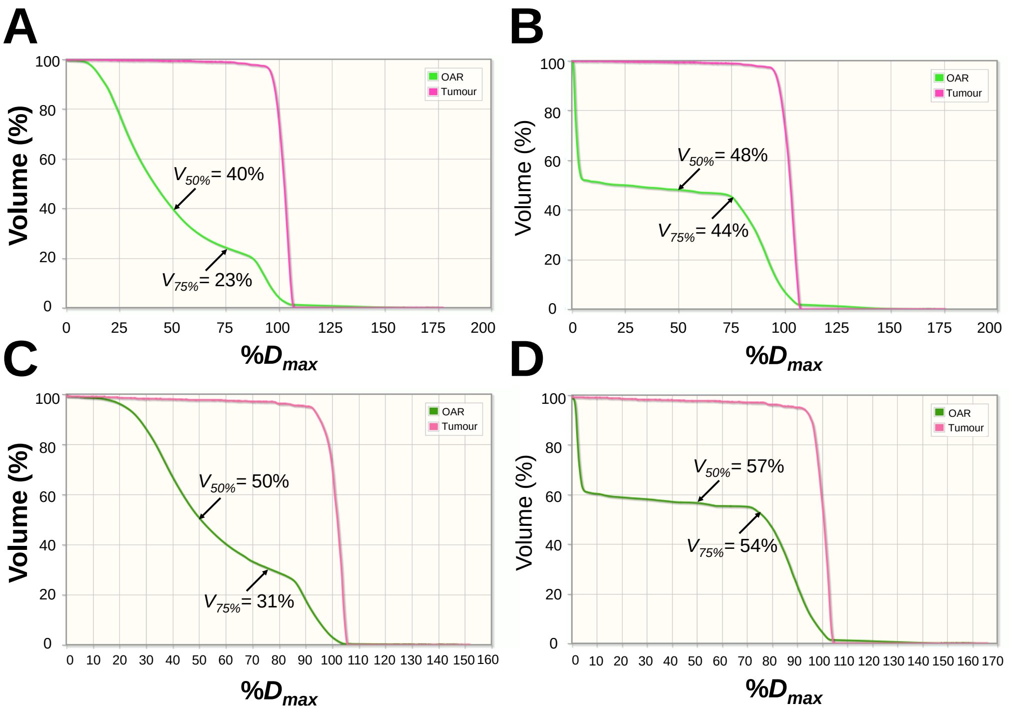


**Supplementary Figure 3. Dose-volume histograms (DVH) for two treatment scenarios in two breast cancer xenografts** with the A,C) moving arcs or B,D) fixed beam. Dosimetric characteristics of target volumes (pink) and organs at risk (green) treated depict that both moving arcs and fixed beam achieved similar target coverage. The former achieved better organs at risk sparing with a mean healthy volume receiving 75% of the dose lower in the two examplar mice.

**Supplementary Table 2. Differences between preclinical image-guided irradiators and conventional clinical linacs, and between dosimetric characteristics of output ionising photon radiation.** *Note: MFP, electron mean free path;* $D_{max}$*, maximal dose in percent depth dose graphs; LET, linear energy transfer;* $Z$*, atomic number; CT, computed tomography; PTV, planned target volume; OAR, organ at risk.*

| | **Preclinical RT** | **Clinical RT** |
|---|---|---|
| **Tube acceleration potential (maximum photon energy**) | 220 kV (220 keV) | 6 MV (6 MeV) or 10 MV (10 MeV) |
| **Skin/surface relative dose** | High (about 100% of $D_{max}$ at skin) | Low (about 40% of $D_{max}$ at skin) |
| **Dose relative fall-off** | Fast (about 40% of $D_{max}$ at 5 cm depth) | Slow (about 80% of $D_{max}$ at 5 cm depth) |
| **Secondary electrons range** | Short (MFP $\approx$ 1-2 mm) | Long (MFP $\approx$ 10-20 mm) |
| **Linear energy transfer (LET)** | Intermediate (LET $\approx$ 3-4 keV/µm) | Low (LET $\approx$ 0.2 keV/µm) |
| **Dominant photon interaction**<br>(Figure **??**B) | *Photoelectric Effect*<br>-All photon energy is transferred in medium and electrons are emitted<br>-Energies <50 keV<br>-Mass absorption coefficient proportional to $Z^3$<br>-Strong backscattering | *Compton Scattering*<br>-Photon transfers some energy to an outer shell electron which is ejected, and then scatters<br>-Energies in 100 keV-10 MeV range<br>-Mass absorption coefficient independent of $Z$<br>-Forward directed |
| **Imaging modality for treatment planning** | *Cone-beam CT*<br>-X-ray tube voltage = 60 kVp<br>-X-ray tube current = 13 mA<br>-Low contrast, prone to image artefacts and fixed resolution<br>-Cone-beam multi-slice simultaneous tomographic acquisition | *CT simulator*<br>-X-ray tube voltage = 70-140 kVp<br>-X-ray tube current = 50-800 mA with automated exposure control<br>-Reconstruction kernel adapted to organ site and desired contrast/resolution<br>-Slice-by-slice tomographic acquisition |
| **Target delineation for treatment planning** | Tumour is targeted with or without PTV definition (not enough contrast to segment) | PTV and OARs are segmented and plan is optimised to limit dose to OARs |

**Supplementary Table 3.** Detailed summary of animals included in the longitudinal preclinical radiation therapy trial. Enrolled mice engrafted with either investigated breast cancer cell lines and their imaging data inclusion or exclusion across acquisition time-points are reported. Colour-coding represent data acquired and included (green), acquired and excluded (yellow) and not acquired (red). Stars represent gas challenge data acquisition during tomographic photoacoustic imaging examination. A total of 11 mice out of 46 had at least one reported exclusion.

| Tumour type | Tx arm | Tumour number | Study ID | IHC | Mesoscopic PAI | | | Tomographic PAI | | | Exclusion & Reason | DoD, days post-RT |
|---|---|---|---|---|---|---|---|---|---|---|---|---|
| | | | | | Pre-RT | Post-RT | Endpoint | Pre-RT | Post-RT | Endpoint | | |
| MCF7 | SDRT | 1 | PAIinRT015 | | | | | * | * | * | | 14 |
| | | 2 | PAIinRT016 | | | | | * | * | * | | 14 |
| | | 3 | PAIinRT017 | | | | | * | * | | Weight loss | 6 |
| | | 4 | PAIinRT032 | | | | | | | * | | 7 |
| | | 5 | PAIinRT033 | | | | | | | * | | 7 |
| | | 6 | PAIinRT034 | | | | | | | * | | 7 |
| | | 7 | PAIinRT047 | | | | | * | * | * | | 7 |
| | | 8 | PAIinRT048 | | | | | * | * | * | | 7 |
| | HFRT | 9 | PAIinRT019 | | | | | * | * | * | | 14 |
| | | 10 | PAIinRT020 | | | | | * | * | * | | 9 |
| | | 11 | PAIinRT025 | | | | | * | * | * | | 7 |
| | | 12 | PAIinRT026 | | | | | * | Excl. | | Anaesthesia, gas challenge | 1 |
| | | 13 | PAIinRT035 | | | | | | | * | | 7 |
| | | 14 | PAIinRT036 | | | | | | | * | | 7 |
| | | 15 | PAIinRT049 | | | | | * | * | * | | 7 |
| | | 16 | PAIinRT050 | | | | | * | * | * | | 7 |
| | | 17 | PAIinRT051 | | | | | * | * | * | | 7 |
| | Control | 18 | PAIinRT039 | | | | | | | * | | 7 |
| | | 19 | PAIinRT040 | | | | | | | * | | 7 |
| | | 20 | PAIinRT052 | | | | | * | * | * | | 7 |
| | | 21 | PAIinRT053 | | | | | * | * | * | | 7 |
| | | 22 | PAIinRT054 | | | | | * | * | * | | 7 |
| | | 23 | PAIinRT055 | | | | | * | * | * | | 7 |
| MDA-MB-231 | SDRT | 1 | PAIinRT009 | | | | | * | * | | Inactive, poor body condition | 6 |
| | | 2 | PAIinRT010 | | | | | * | * | * | | 14 |
| | | 3 | PAIinRT018 | | | | | * | * | | Weight loss | 5 |
| | | 4 | PAIinRT022 | | | | | * | * | * | | 8 |
| | | 5 | PAIinRT023 | | | | | * | * | | Inactive, poor body condition | 6 |
| | | 6 | PAIinRT024 | | | | | * | * | | Weight loss | 6 |
| | | 7 | PAIinRT027 | | | | | * | * | | Anaesthesia | 7 |
| | | 8 | PAIinRT028 | | | | | * | * | * | | 7 |
| | | 9 | PAIinRT030 | | | | | | | * | Early endpoint due to poor body condition | 4 |
| | | 10 | PAIinRT031 | | | | | | | * | Early endpoint due to poor body condition | 4 |
| | HFRT | 11 | PAIinRT012 | | | | | * | * | * | | 14 |
| | | 12 | PAIinRT013 | | | | | * | * | | Weight loss | 2 |
| | | 13 | PAIinRT014 | | | | | * | * | | Weight loss | 2 |
| | | 14 | PAIinRT021 | | | | | * | * | * | Ulceration | 9 |
| | | 15 | PAIinRT037 | | | | | | | * | | 7 |
| | | 16 | PAIinRT038 | | | | | | | * | | 7 |
| | Control | 17 | PAIinRT029 | | | | | * | * | * | | 7 |
| | | 18 | PAIinRT041 | | | | | | | * | | 7 |
| | | 19 | PAIinRT042 | | | | | * | * | * | | 7 |
| | | 20 | PAIinRT043 | | | | | * | * | * | | 7 |
| | | 21 | PAIinRT044 | | | | | * | * | * | | 7 |
| | | 22 | PAIinRT045 | | | | | * | * | | Anaesthesia | 7 |
| | | 23 | PAIinRT046 | | Excl. | Excl. | Excl. | * | * | * | Significant ulceration | 7 |

**Supplementary Table 4.** Summary table of extracted quantitative photoacoustic imaging biomarkers in both modalities. Note: $i$, pixel element; $N$, total pixel within region of interest.

| Modality | Quantitative imaging biomarker | Physical quantity measured | Biological processes represented |
|---|---|---|---|
| Multispectral tomographic photoacoustic imaging | Total haemoglobin (THb), sum of deoxy-haemoglobin (HbR) and oxyhaemoglobin ($HbO_2$) | THb = $\sum_{i=1}^{N}(\mathrm{HbR}_i + \mathrm{HbO}_{2i})/N$ | Blood content |
| | Blood oxygen saturation ($sO_2$) | $sO_2 = \sum_{i=1}^{N}(\mathrm{HbO}_{2i}\ /\ (\mathrm{HbR}_i + \mathrm{HbO}_{2i}))/N$ | Tissue blood oxygenation |
| | Standard deviation (SD) of $sO_2$ | SD of $sO_2 = \sqrt{\sum_{i=1}^{N}(\mathrm{sO}_{2i} - \mathrm{sO}_2)^2/N}$ | Intratumoural blood oxygen heterogeneity |
| | Change in $sO_2$ under gas challenge ($\Delta sO_2$) and responding fraction (RF) | $\Delta sO_2 = sO_2^{100\%O_2} - sO_2^{Air}$; RF defined as pixel count with $\Delta sO_2$ >mean + 1 SD of $\Delta sO_2$ over total number of pixels | Tissue oxygen diffusion |
| Monospectral mesoscopic photoacoustic imaging | Blood volume (BV) | Total pixels in segmented vasculature | Superficial perfused tumour blood content |
| | Diameter | Average segments diameter in segmented skeletonised vascular network | Diameter of vessels across the tumour periphery |
| | Loop 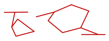 | Count of groups of vertices connected by path forming a closed structure in segmented skeletonised vascular network | Looping or curving structures in the vasculature at the tumour periphery |
| | Connected component (CC) 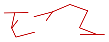 | Count of groups of vertices connected by path with edges in segmented skeletonised vascular network | Connectivity of the vasculature |
| | Density | Counts of vessel segments normalised to BV | Vessel coverage in tumour periphery |

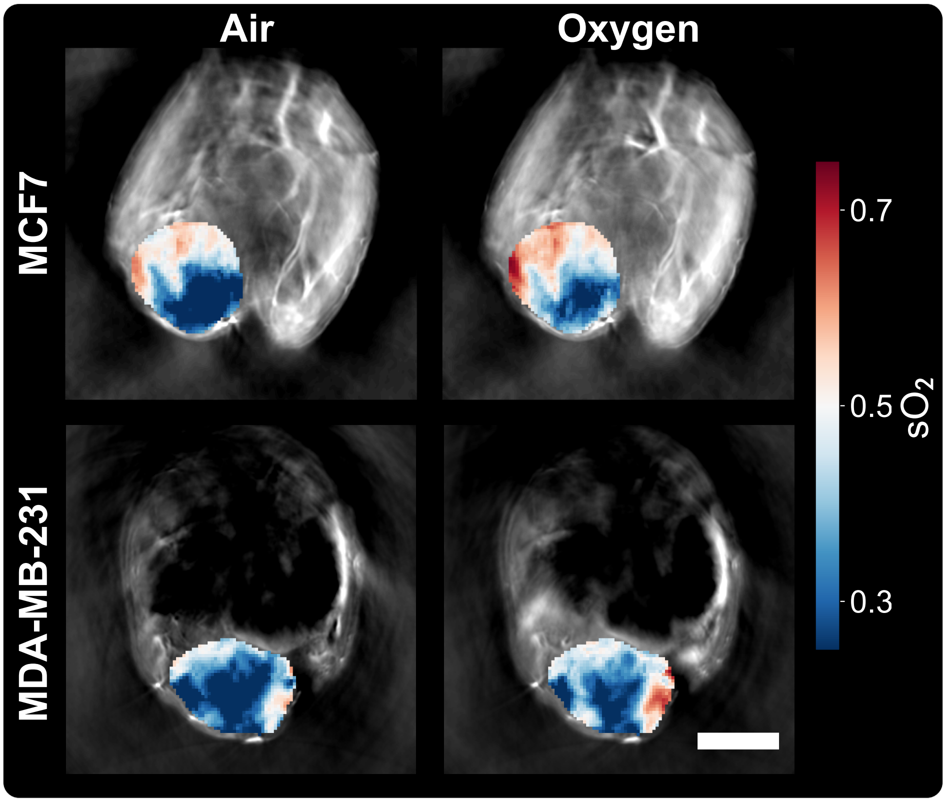


**Supplementary Figure 4. Exemplar MCF7 and MDA-MB-231 tumour-bearing mice at baseline imaged with multispectral tomographic PAI with quantitative parametric maps of blood oxygen saturation ($sO_2$) under gas challenge.** Tumour $sO_2$ displays higher deoxyhaemoglobin content (blue) under breathing air (20& $O_2$), and higher oxyhaemoglobin content (red) when breathing gas is switched to pure oxygen delivery. Calculation of the pixel-wise difference between the two parametric maps allow the quantification of the change in $sO_2$ and the responding fraction. Scale bar, 2 mm

**Supplementary Table 5.** Immunohistochemistry and histopathological markers used for staining processed tumour sections and quantification.

| Antibody | Supplier | Retrieval method | Visualisation | Quantification |
|---|---|---|---|---|
| Anti-mouse Cluster of Differentiation 31 (CD31) | Cell Signaling, 77699 | 1:100, Tris-EDTA HIER 20min | Stable and highly expressed endothelial cell marker | Percent CD31 positive area in non-necrotic viable tumour classified with trained random forest classifier |
| Anti-mouse Alpha Smooth Muscle Actin (ASMA) | Abcam, ab5694 | 1:500, Tris-EDTA HIER 10min | Smooth muscle and pericyte marker; indicates vascular maturity | Percent ASMA positive area in non-necrotic viable tumour classified with trained random forest classifier; overlaid with CD31-classified area for capturing double CD31-ASMA area |
| Anti-human Ki67 | Daco, Agilent, M7240 | 1:400, Tris-EDTA HIER 30min | Protein expressed in active phases of cellular division, marker of proliferation | Percent Ki67 positive nuclei over all nuclei in non-necrotic viable tumour classified with trained random forest classifier |
| Anti-human phosphorylated H2A Histone Family Member X ($\gamma$-H2AX) | Cell Signaling, 9718 | 1:200, Sodium Citrate HIER 20min | Core protein that structures DNA into chromatine and becomes phosphorylated with strand breaks, marker of DNA damage | Percent $\gamma$-H2AX positive nuclei over all nuclei in non-necrotic viable tumour classified with trained random forest classifier |
| Anti-human Hypoxia inducible factor 1-$\alpha$ (HIF1-$\alpha$) | Abcam, ab51608 | 23.36 µg/ml, Sodium Citrate HIER 20min | Acute hypoxia marker | Percent HIF1-$\alpha$ positive nuclei over all nuclei in non-necrotic viable tumour classified with trained random forest classifier |
| Haematoxylin & Eosin (H&E) | Leica Microsystems Eosin 1%, Haem. 3801560E | – | Combined nuclear and cytoplasm/extracellular tissue markers | Percent necrotic area classified across the whole tumour, excluding skin regions for all analyses, with trained random forest classifier |

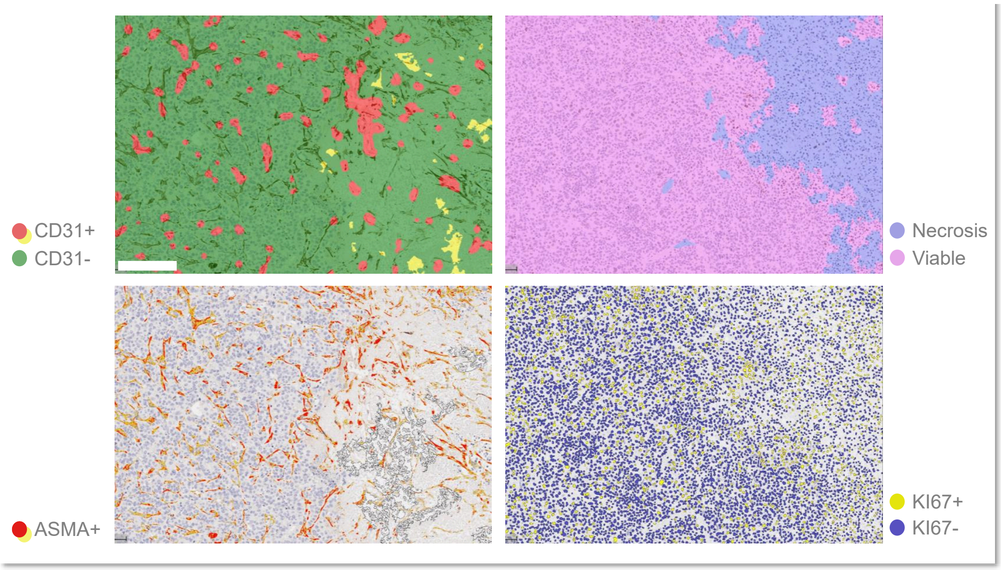


**Supplementary Figure 5. Immunohistochemistry (IHC) analysis of resected tumour tissue sections from breast cancer xenograft.** ASMA stained IHC panel (left) with overlaid CD31 classifier (top) trained using HALO showing areas classified as positive (red) and negative (green), and with overlaid ASMA classifier (bottom) showing positive areas (strongly positive in red). Ki67 sequential stained IHC section panel with overlaid necrosis (purple) vs. viable tissue (pink) classifier (top) and with overlaid Ki67 positive nuclei (yellow) and negative nuclei (blue).

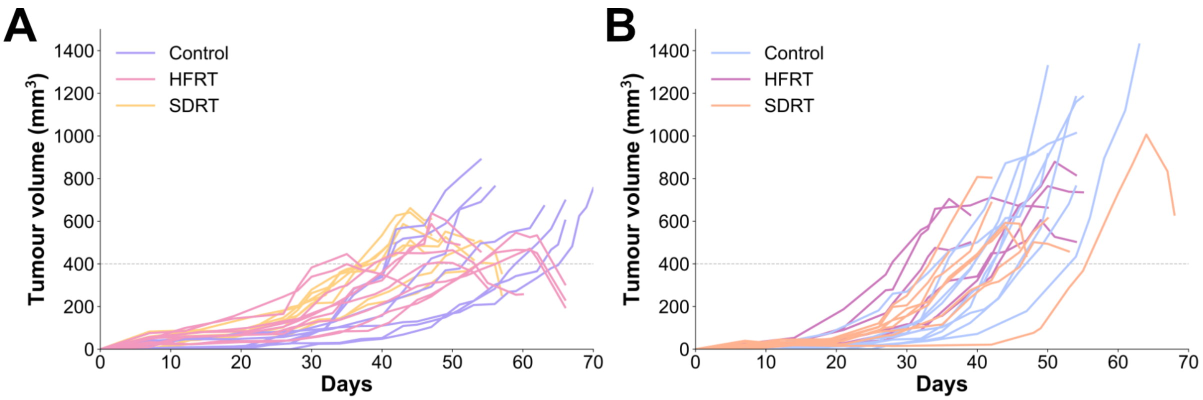


**Supplementary Figure 6. Tumour growth curves in individual mice enrolled in the preclinical radiation therapy trial.** Tumour volumes capture by calliper measurements from inoculation date to endpoint in A) MCF7 and B) MDA-MB-231 tumour-bearing xenograft models.

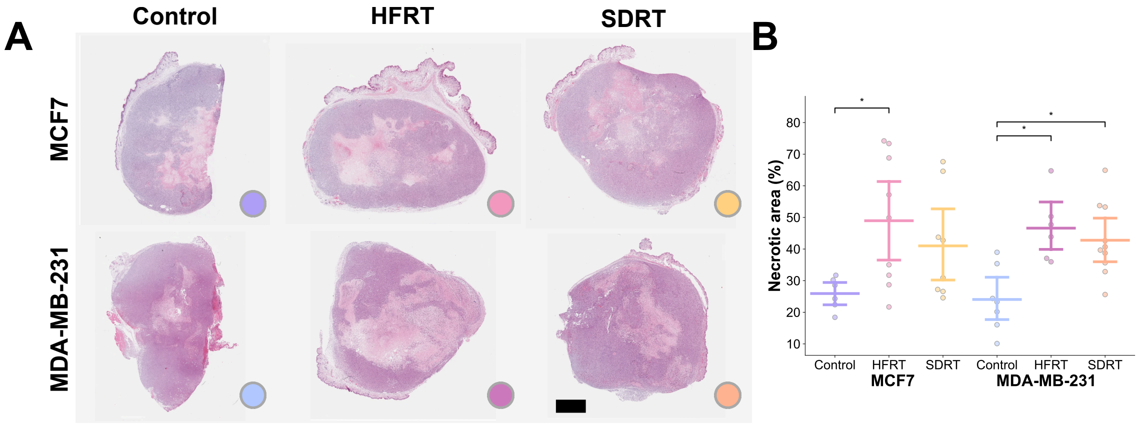


**Supplementary Figure 7. Exemplar haematoxilin and eosin stain scans of resected tumour tissue sections** in A) MCF7 (top panels) and MDA-MB-231 (bottom panels) breast cancer xenografts across treatment conditions. B) Quantified percent necrosis area across models and treatment groups in bar and point plots with bars representing mean and 95% confidence intervals. Scale bar, 1.5 mm.

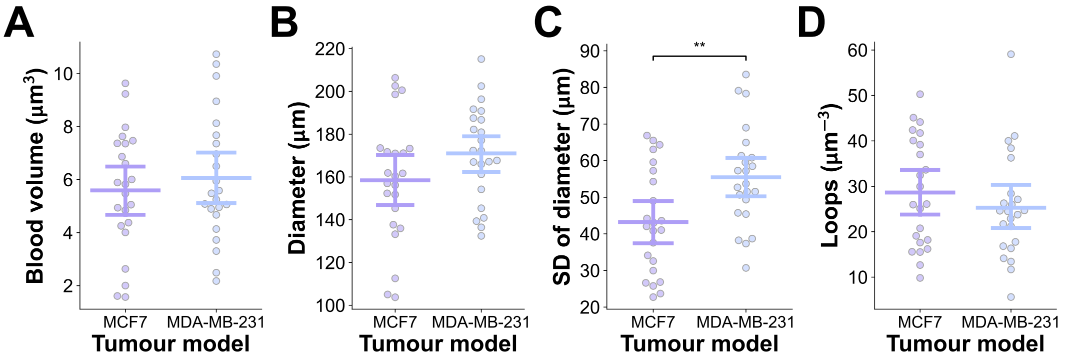


**Supplementary Figure 8. Baseline mesoscopic photoacoustic imaging parameters quantified in segmented vascular networks.** MCF7 and MDA-MB-231 xenografts' vasculature quantified for A) blood volume, B) average diameter, C) standard deviation of diameter, and D) loops normalised to blood volume in bar and point plots, with bars representing mean and 95% confidence intervals.

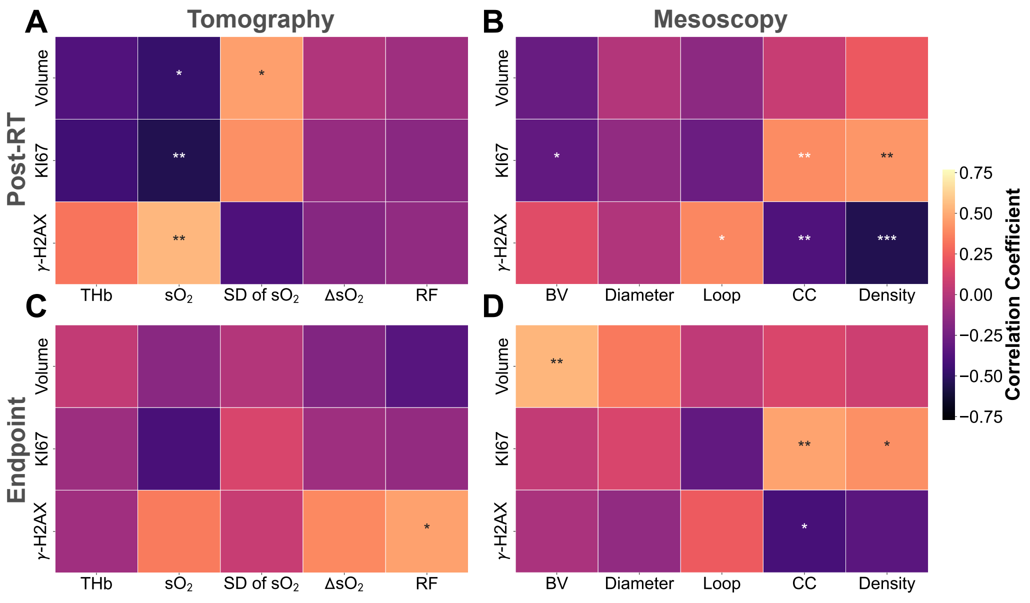


**Supplementary Figure 9. Bivariate correlation of quantitative photoacoustic imaging biomarkers with endpoint immunohistochemistry.** Correlation heatmaps of post-RT A) tomographic and B) mesoscopic PAI biomarkers, and of endpoint C) tomographic and D) mesoscopic PAI biomarkers with *ex vivo* immunohistochemistry parameters.
